# Multispectral Fingerprinting Resolves Dynamics of Nanomaterial Trafficking in Primary Endothelial Cells

**DOI:** 10.1101/2020.12.14.422763

**Authors:** Mitchell Gravely, Daniel Roxbury

## Abstract

Intracellular vesicle trafficking involves a complex series of biological pathways used to sort, recycle, and degrade extracellular components, including engineered nanomaterials which gain cellular entry *via* active endocytic processes. A recent emphasis on routes of nanomaterial uptake has established key physicochemical properties which direct certain mechanisms, yet relatively few studies have identified their effect on intracellular trafficking processes past entry and initial subcellular localization. Here, we developed and applied an approach where single-walled carbon nanotubes (SWCNTs) play a dual role - that of an engineered nanomaterial (ENM) undergoing intracellular processing, in addition to functioning as the signal transduction element reporting these events in individual cells with single organelle resolution. We used the unique optical properties exhibited by non-covalent hybrids of single-stranded DNA and SWCNTs (DNA-SWCNTs) to report the progression of intracellular processing events *via* two orthogonal hyperspectral imaging approaches of near-infrared (NIR) fluorescence and resonance Raman scattering. A positive correlation between fluorescence and G-band intensities was uncovered within single cells, while exciton energy transfer and eventual aggregation of DNA-SWCNTs were observed to scale with increasing time after internalization. These were confirmed to be consequences of intracellular processes using pharmacological inhibitors of endosomal maturation, which suppressed spectral changes through two distinct mechanisms. An analysis pipeline was developed to colocalize and deconvolute the fluorescence and Raman spectra of subcellular regions of interest (ROIs), allowing for single-chirality component spectra to be obtained with sub-micron spatial resolution. This approach uncovered a complex relationship between DNA-SWCNT concentration, fluorescence intensity, environmental transformations, and irreversible aggregation resulting from the temporal evolution of trafficking events. Finally, a spectral clustering analysis was applied to delineate the dynamic sequence of processes into four distinct populations, allowing stages of the intracellular trafficking process to be identified by the multispectral fingerprint of encapsulated DNA-SWCNTs.

**TOC Graphic:** 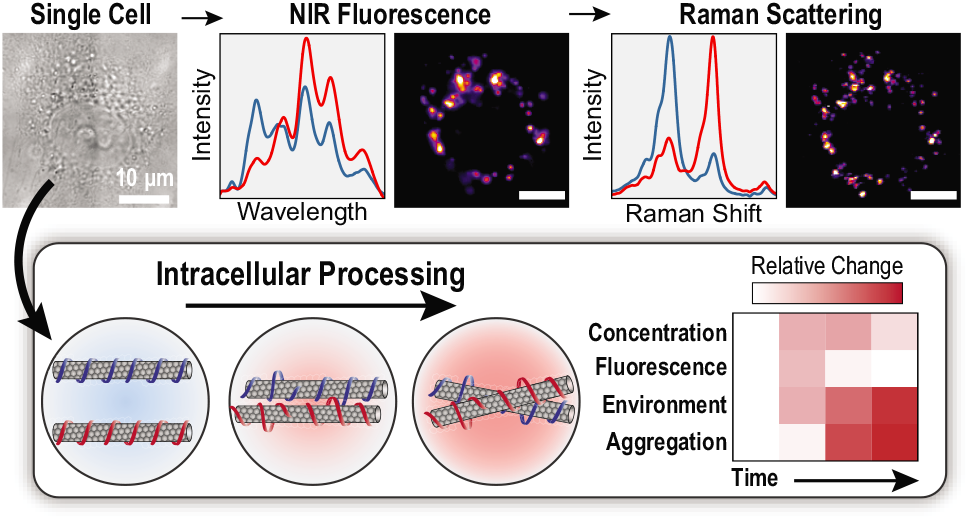

## Introduction

Intracellular trafficking is a highly regulated yet diverse system of pathways involving the entry, translocation, and localization of cargo internalized by endocytic cells.^1–3^ The endosomal maturation process, which initializes the main cellular degradation pathway, entails a dramatic series of physicochemical changes; a drop in luminal pH, influx of lysosomal enzymes, and change of ionic environment all promote digestion of vesicle contents.^3^ Most types of engineered nanomaterials (ENMs) gain cellular entry through active endocytic processes,^4^ where they are trafficked through these endosomal pathways before accumulating in lysosomal vesicles.^5, 6^ The key mechanisms of entry and localization of ENMs in biological systems have been extensively studied.^7^ The differential uptake of ENMs based on their size,^8^ shape,^9^ and surface chemistry^10^ have provided insight on targeting, while the formation of a protein corona on the ENM surface is a dynamic process that can further direct biological interactions.^11, 12^ In contrast, the pathways in which these ENMs are subject to after internalization, as well as the effects they might have on these pathways, lack the same depth of clarity despite the importance of these native processes. Proper lysosomal function and trafficking are essential in multiple metabolic pathways which regulate basic cellular functions including autophagy, nutrient degradation, and catabolite export.^6^ In addition, dysfunction of the endosomal-lysosomal pathways has been implicated in Alzheimer’s disease,^13^ lysosomal storage disorders,^14, 15^ and infectious diseases.^16^ As EMNs are developed for biomedical applications, it is crucial to understand their environmental interactions in complex biological systems at the single-organelle level in order to properly assess their impact on key cellular functions. Intracellular trafficking is accurately described by a range of characteristics across a population of vesicles due to the asynchronous nature of the endocytic pathway,^3^ and thus cell-averaged analysis leads to systematic and compound errors. By tracking the fate of individual endosomal pathway reporters, a more accurate representation of the distribution of processes can be obtained to provide a deeper understanding of the dynamic ENM trafficking system.

The continuous evolution of endocytic vesicles presents a challenging system to investigate since experimental strategies are often limited by the optical capabilities of a given ENM. Fluorescence microscopy and organelle colocalization have routinely been used, providing valuable insight on the spatial and temporal localization of internalized ENMs,^17, 18^ however traditional fluorophores lack environmental responsivity. Hyperspectral microscopy and confocal Raman imaging, which can both resolve spectral data with spatial resolution, are two approaches that can provide information about physical and chemical components within a system. Nearinfrared (NIR) hyperspectral fluorescence imaging has enabled environmental sensing within endosomal vesicles of live cells,^5, 19, 20^ while Raman probes have been designed to report intracellular aggregation,^21^ pH,^22^ as well as the molecular composition within various endosomal vesicles.^23^ These approaches provide robust intracellular data which can characterize the complex ENM interactions in biological settings, however more suitable methods for processing and interpreting these highly dimensional datasets must be developed to enable widespread adoption of these techniques.^24^

Single-walled carbon nanotubes (SWCNTs) are among the distinctive materials which exhibit both NIR fluorescence and resonance Raman scattering as intrinsic optical properties,^25^ making them exceptional reporters of physical and environmental changes. The electronic structure of a given SWCNT, including its metallic or semiconducting character,^26^ is dependent on its chiral identity (denoted by the integers (*n,m*)), which varies by diameter and roll-up angle. Each chirality possesses unique optical transition energies between valence and conduction bands (*E_ii_*, where *i* = 1, 2, etc.),^27^ and as a result, the intensity of the Raman spectrum from SWCNTs with *E_ii_* resonant with the laser excitation is significantly enhanced by resonance Raman scattering.^28^ At the same time, semiconducting SWCNTs exhibit band gap fluorescence when excited at their *E_22_* resonances (500-900 nm),^29^ however the two observed spectra provide unique information which can detail their physical state and local environment. The Raman spectrum contains multiple features, most notably the radial breathing mode (RBM, 150 – 350 cm^−1^) and G-band (~1589 cm^−1^), which can be used to characterize the chiral composition,^27^ concentration,^30^ aggregation state,^31^ and surface chemistry^32^ of a SWCNT mixture. SWCNTs emit fluorescence in the NIR range (~900 – 1400 nm), where absorbance and scattering effects from biological samples are at a minimum,^33^ to produce a multi-peak spectrum of all excitable chiralities at given excitation wavelength. Because SWCNTs exhibit solvatochromism,^34^ the emission from each chirality is subject to position and intensity modulation in response to environmental changes, including analyte binding,^35^ changes in charge density,^36^ aggregation,^37^ pH,^29^ and ionic environment.^38^

Single-stranded DNA, which can disperse single SWCNTs into a stable aqueous suspension,^39^ provides a biocompatible surface functionalization while preserving their advantageous optical properties.^40, 41^ DNA-SWCNTs are internalized by cells *via* energy dependent endocytosis and localize to intracellular vesicles in the endolysosomal pathway,^5, 19^ making them exceptional candidates to nonspecifically target these trafficking processes. Therefore, we propose that simultaneous characterization of (1) the intracellular environmental conditions and (2) the ENMs physical condition can be achieved using DNA-SWCNTs, allowing for a multispectral characterization of the intracellular trafficking processes. Here, we report the internalization and intracellular processing of DNA-SWCNTs within individual cells using a dual-hyperspectral colocalization technique, which correlated intracellular fluorescence and Raman spectra. In tandem, the responsive photoluminescence and multi-featured Raman scattering from

DNA-SWCNTs detail the changing intracellular environment and the resultant condition of the SWCNT hybrids in primary endothelial cells. We observed a temporal increase in local concentration of DNA-SWCNTs, inducing exciton energy transfer (EET)^42^ and aggregation at two distinct rates within concentrated regions due to vesicle coalescence during intracellular processing. Pharmacological inhibitors of endosomal maturation effectively eliminated these events, confirming these processes were responsible, while a DNA sequence dependence was generally not observed. Common regions of interests (ROIs) were identified within individual cells to colocalize the subcellular regions and spectral deconvolution was performed to obtain single-chirality component spectra. A relationship between concentration, aggregation, and photoluminescence modulation was identified within nanoscale regions, exhibiting heterogeneity which varied in time. Finally, a spectral clustering analysis identified four distinct types of ROIs which mapped the trafficking process, allowing individual vesicles to be classified by their spectral fingerprint without temporal input. To our knowledge, this process details the first time that single chirality SWCNT resolution was achieved simultaneously from Raman and fluorescence spectra acquired within single cells, providing a viable framework for unparalleled intracellular ENM characterization.

## Results and Discussion

### Co-dependence of Fluorescence and G-band Intensities

To develop a spectral model of nanomaterial trafficking, we first identified two formulations of DNA-SWCNTs which were previously shown to induce differential cellular responses when internalized by macrophages as potential intracellular reporters.^20^ HiPco SWCNTs were aqueously dispersed with (GT)_6_, or (GT)_30_ oligonucleotides *via* probe-tip sonication and high-speed ultracentrifugation, resulting in highly purified, monodisperse DNA-SWCNT suspensions.^43^ The presence of multiple peaks in both the visible and NIR range of their absorbance spectrum (Fig. S1a) confirmed that both ssDNA sequences had suspended a multi-chiral mixture with strong optical absorbance. Excitation with a 730 nm laser produced bright fluorescence in the NIR range from multiple chiralities (Fig. S1b), while the apparent differences in peak emission wavelengths were explained by the DNA sequence and nanotube chirality dependence on the hybrid structures.^44^ A 1.58 eV (785 nm) laser source was used to acquire the Raman spectrum of both DNA-SWCNTs (Fig. S1c,d), producing sharp peaks in both the radial breathing mode range (RBM, 150 – 350 cm^−1^) and the G-band (~1589 cm^−1^). Furthermore, the low intensity of the D-band (~1350 cm^−1^) from both samples confirmed the removal of catalyst impurities and amorphous carbon from the raw HiPco materials.^45^

Human umbilical vein endothelial cells (HUVEC primary cell line), a common *in vitro* model used to study neovascularization,^46^ were chosen to represent the endothelium, which would contact any ENMs delivered through intravenous injection. First, HUVEC cultured in grid-labeled glass bottom petri dishes were incubated with 1 mg-L^−1^ of either (GT)_6_- or (GT)_30_-SWCNTs for 1 hour under standard cell culture conditions, after which the SWCNT-containing media was removed and the cells were rinsed with phosphate buffered saline (PBS). Next, the cells were either fixed with paraformaldehyde (considered the 0 hour (h) time point) or replenished with fresh media and allowed to incubate for additional time before fixation. Multiple cells from each condition, identifiable by their location within the grid-labeled culture area, were then imaged at 100 × magnification using both NIR hyperspectral fluorescence and confocal Raman microscopes. Near identical images of the internalized DNA-SWCNTs were constructed from the broadband NIR fluorescence and G-band spectral regions (Fig. 1a-c), each of which depicted distinct subcellular regions containing the (GT)_30_-SWCNTs. Histograms of the NIR fluorescence (Fig. 1d) and G-band intensities (Fig. 1e) were constructed using pixel values from the entire dataset, revealing common temporal changes in the intensity distributions. To quantify these trends, the average intensity fold changes with respect to 0h averages were computed (Fig. 1f), showing nearly identical increases of fluorescence and G-band intensities at 6h followed by differential reductions from 6-24h. The G-band intensity, which is linearly dependent on SWCNT concentration,^30^ could only increase over 6h due to localized concentration increases in the cell, which we hypothesized could be due to fusion of intracellular vesicles over time. Moreover, we suspect the reduced fluorescence at 24h could indicate an intracellular quenching mechanism such as DNA-SWCNT aggregation. Similar results were obtained from cells incubated with (GT)_6_-SWCNTs throughout this study, indicating no clear dependence on DNA sequence. These results can be found in supporting information.

**Figure 1:**
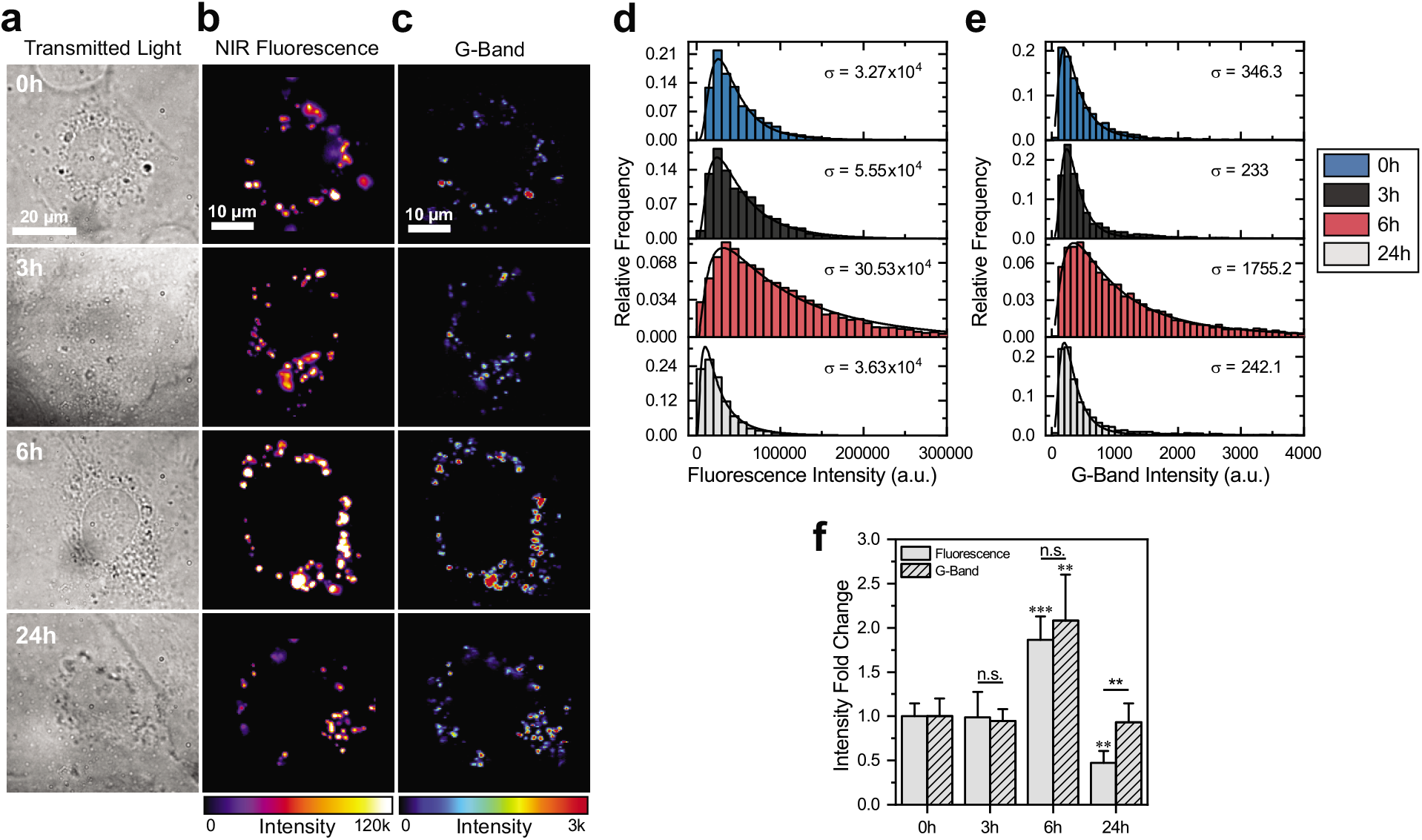
Fluorescence intensity and local concentration of DNA-SWCNTs are co-dependent within single cells. **(a)** Transmitted light, **(b)** broadband NIR fluorescence (950-1350 nm), and **(c)** G-band Raman intensity micrographs of individual cells dosed with 1 mg-L^−1^ (GT)_30_-SWCNTs for 1h and incubated in fresh media for indicated times. **(d)** Fluorescence intensity and **(e)** G-band intensity histograms of SWCNT-containing pixels from all examined cells at each time point. The distributions are fitted to log-normal curves and the widths are estimated by the log standard deviation parameter (σ). **(f)** Fold change of average fluorescence and G-band intensities with respect to 0h averages. Error bars represent mean ± s.d. with *n* ≥ 4 cells per condition. Five pointed stars between columns represent significance between fluorescence and G-band intensities and six pointed stars above columns represent significance versus 0h values. (\**p* < 0.05, \*\**p* < 0.01, \*\*\**p* < 0.001 according to two-tailed two-sample t-test).

### Spectral Features Identify Intracellular Aggregation of DNA-SWCNTs

While the broadband fluorescence intensity from DNA-SWCNTs is the integrated sum of all excited chiralities, the intensity from each individual chirality can be differentially affected.^44^ By splitting the fluorescence spectrum from internalized DNA-SWCNTs into emission wavelength bands (Fig. 2a), the integrated intensities corresponding to chiralities emitting over various wavelength windows could be quantified independently (Table S1).^36^ The integrated intensity from each band was normalized by the total intensity from each cell and the average normalized intensities were computed (Fig. 2b), illustrating the relative intensity change of each band over time. We found that bands with lower emission energies (higher wavelength) generally increased over time, while the intensities of the two highest energy bands either decreased or remained constant, revealing certain chirality dependences. The ratiometric intensity of band 4 divided by band 1 (Fig. 2c) provided a metric of this trend, which could be fitted to an exponential curve with respect to time. The decreasing and increasing of high and low energy emission intensities, respectively, are characteristics observed from exciton energy transfer (EET) between individual SWCNT chiralities in close proximity,^42, 47^ indicating a progressive degree of DNA-SWCNT flocculation occurring in time due to intracellular processing.

**Figure 2:**
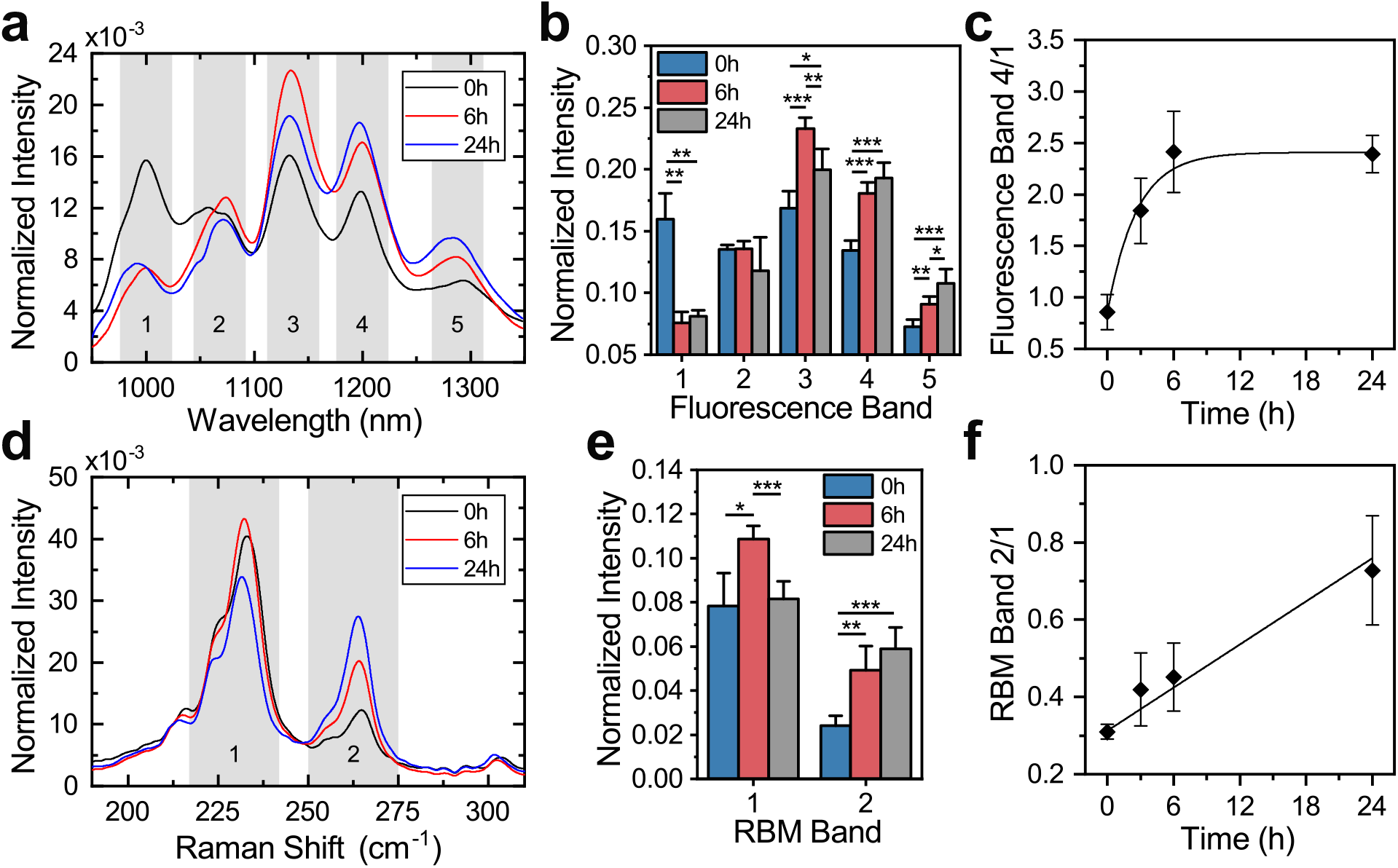
Temporal resolution of DNA-SWCNT spectral features indicates aggregation within subcellular regions. **(a)** Average fluorescence spectrum of (GT)_30_-SWCNTs in single cells after variable lengths of intracellular processing, normalized to the total integrated intensity of each spectrum. Fluorescence bands are indicated by shaded regions. **(b)** Average normalized fluorescence band intensities from (GT)_30_-SWCNTs in single cells after variable lengths of intracellular processing. Each spectrum was normalized by the total cell intensity, and average normalized band intensities are reported. **(c)** Ratiometric intensity of fluorescence band 4 divided by band 1, with exponential fit, as a function of time. **(d)** RBM region of the average Raman spectrum of (GT)_30_-SWCNTs in single cells after variable lengths of intracellular processing, normalized to the total integrated intensity of each spectrum. RBM bands are indicated by shaded regions. **(e)** Average normalized RBM band intensities from (GT)_30_-SWCNTs in single cells after variable lengths of intracellular processing. Each spectrum was normalized by the total cell RBM intensity, and average normalized band intensities are reported. **(f)** Ratiometric intensity of RBM band 2 divided by band 1, with linear fit, as a function of time. Error bars represent mean ± s.d. for all, with *n* ≥ 4 cells per condition. (\**p* < 0.05, \*\**p* < 0.01, \*\*\**p* < 0.001 according to two-tailed two-sample t-test).

The same band deconvolution process was applied to the two dominant regions of the RBM spectrum (Fig. 2d), and the average normalized intensity from cells at each time point were determined (Fig. 2e), revealing a monotonic increase of band 2 with increasing incubation time. The ratiometric intensity of band 2 divided by band 1 (Fig. 2f) quantified the intensity changes, displaying a linear increase in time. To explain these findings, we acquired Raman spectra of DNA-SWCNTs both in solution and aggregated out of solution (Fig. S3) and identified the contributing chiralities present in each band along with their *E_22_* transition energies when dispersed in a solution (Table S2).^48^ In general, chiralities in band 1 were within the excitation resonance range of the 1.58 eV laser, resulting in higher intensity RBM features compared to band 2 when in solution. However, the dramatic increase of band 2 intensities upon aggregation is explained by a decrease in *E_22_* transition energies, bringing these chiralities into resonance with the laser while simultaneously shifting band 1 chiralities out of resonance.^31^ Therefore, we attribute the increase of band 2 intensity over time to intracellular aggregation of DNA-SWCNTs, in agreement with previous findings,^21^ and propose the ratiometric intensity of band 2 divided by band 1 could be used to quantify the degree of aggregation. The fluorescence and RBM band ratios established could potentially discern between tightly compacted and irreversibly aggregated DNA-SWCNTs, respectively, due to their differing responses to complete SWCNT bundling. The fluorescence intensity of DNA-SWCNTs rapidly decreases upon formation of hard aggregates (i.e., direct contact between exposed SWCNT surfaces),^49, 50^ eventually causing EET to reach a maximum level before becoming undetectable due to fluorescence quenching. In contrast, the RBM remains optically active regardless of dispersion quality, and thus a transition from closely packed DNA-SWCNTs to directly aggregated SWCNT bundles could be identified as the point where fluorescence band 4/1 plateaus and RBM band 2/1 continues to increase.

### Inhibition of Endosomal Maturation Reduces Spectral Changes

To confirm the observed spectral changes were induced *via* vesicle trafficking and endosomal maturation, we investigated the effect of inhibiting these native processes using two mechanistically different pharmacological inhibitors. HUVEC cells were incubated with DNA-SWCNTs following the same procedure previously described, however the cells were treated with 10 μg-mL^−1^ Nocodazole (NOC), which polymerizes microtubules and inhibits vesicle motility,^51^ or 100 μM Chloroquine (CQ), which elevates the luminal pH of endosomal vesicles,^52^ for 6h following DNA-SWCNT removal. The fold change of G-band and fluorescence intensities with respect to 0h averages were computed for cells treated with both compounds (Fig. 3a,b), revealing inhibited increases of G-band intensity from both treatments and a reduction of fluorescence from CQ when compared to the 6h control condition. We extended this analysis to examine the effect of pharmacological inhibition on intracellular EET and aggregation by calculating the fluorescence band 4 divided by band 1 intensity ratio (Fig. 3c) and RBM band 2 divided by band 1 intensity ratio (Fig. 3d). Significant inhibition of EET occurred from both treatments, while a high degree of variability in aggregation from individual cells was observed. Notably, the two treatments differentially affected the processes of intracellular trafficking and endosomal maturation, resulting in spectral similarities between 0h or 3h untreated cells and 6h CQ or NOC treated cells, respectively (Fig. 3e,f). This could be explained by their differing mechanisms of action; CQ prevents endosomal maturation and vesicle fusion by directly inhibiting endosomal acidification,^53^ while NOC does not inhibit the initial acidification of endosomes,^54^ but rather prevents cargo from reaching and fusing with more acidic organelles.^51^ This could allow the initial steps of vesicle maturation to occur during treatment with NOC, while initiation of these processes was immediately inhibited following treatment with CQ.

**Figure 3:**
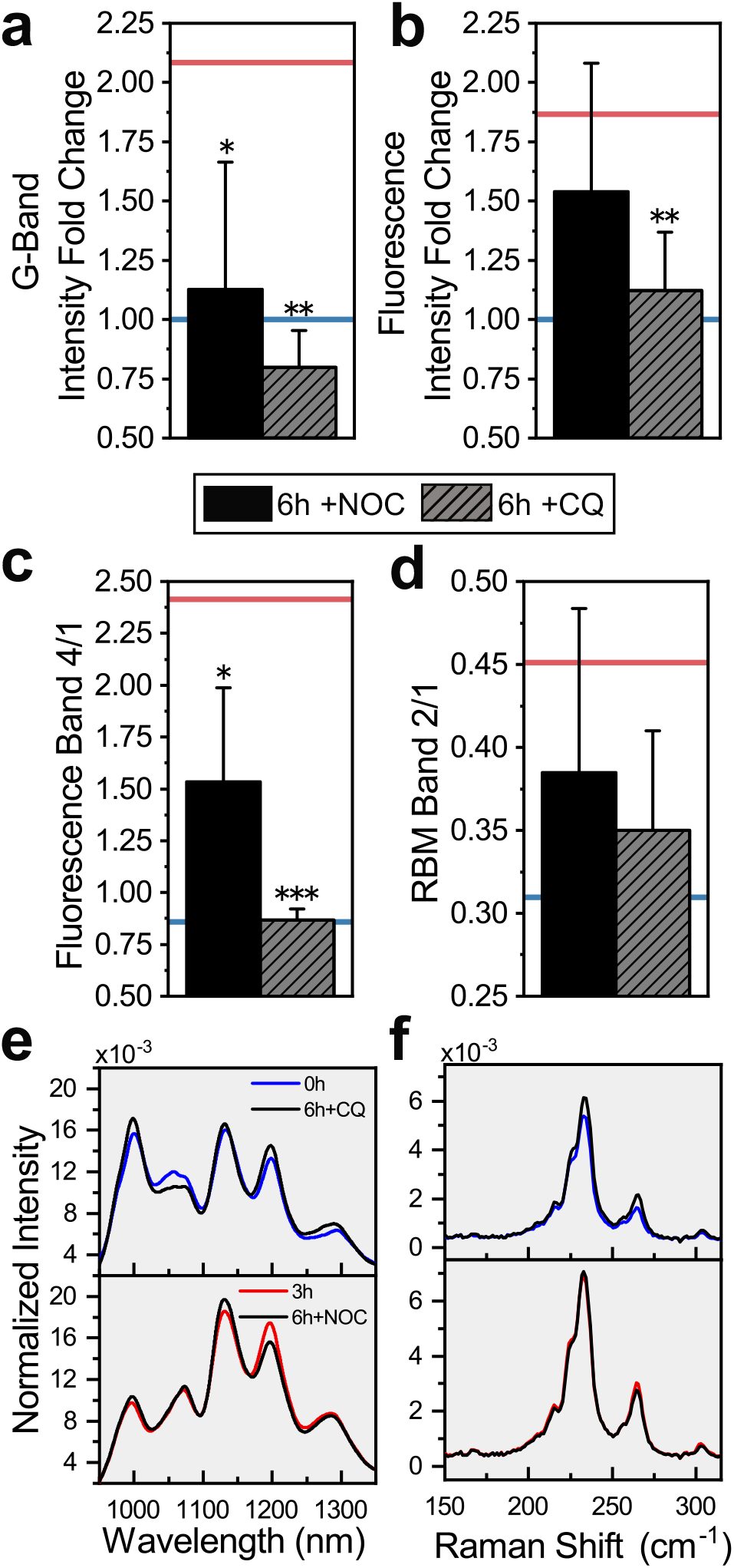
Spectral response to inhibition of endosomal progression. **(a)** Fold change of G-band and **(b)** fluorescence intensities, with respect to 0h controls, from intracellular (GT)_30_-SWCNTs after 6h of incubation with Nocodazole (NOC, 10 μg-mL^−1^) or Chloroquine (CQ, 100 μM). Averages from untreated cells at 0h or 6h are shown as blue or red lines, respectively. **(c)** Ratiometric intensity of fluorescence band 4 divided by band 1 and **(d)** RBM band 2 divided by band 1 from inhibitor-treated cells after 6h. Error bars represent mean ± s.d. for all, with *n* ≥ 4 cells per condition. Stars above error bars represent significance versus 6h untreated cells. (\**p* < 0.05, \*\**p* < 0.01, \*\*\**p* < 0.001 according to two-tailed two-sample t-test). **(e)** Average intracellular fluorescence and **(f)** RBM spectra from inhibitor-treated cells after 6h compared with spectra from untreated cells at indicated times. Each spectrum was normalized to the total integrated intensity.

### Segmentation and Colocalization of Intracellular ROIs

Spectra acquired within whole cells can provide ensemble measurements of internalized DNA-SWCNTs, yet the endocytic system is heterogeneous by nature due to a lack of synchrony between processes occurring simultaneously,^3^ potentially resulting in measurement bias towards more abundant processes while eliminating observation of rare occurrences within single vesicles.^17^ To overcome this issue, we developed a method which could segment a single cell into multiple regions of interest (ROIs) while colocalizing the signals from fluorescence and Raman spectra. First, NIR fluorescence and G-band intensity images (Fig. 4a) were constructed, roughly colocalized, and binarized to create two equally sized template images. The template images were then segmented into separate matching ROIs (Fig. 4b), which were individually confirmed and adjusted manually to account for processing errors and minor discrepancies in ROI locations. The pixels within each ROI were then averaged to create a single fluorescence and Raman spectrum belonging to each subcellular region (Fig. 4c). Finally, the full fluorescence spectrum and the RBM of the Raman spectrum were deconvoluted to their chirality components using simultaneous multipeak fitting algorithms modeled by Voigt^55^ and Lorentz^56^ line shapes, respectively, while the G-band was fit independently to a single Lorentz curve. The fits were restricted by known peak characteristics from literature,^48, 56^ and the whole process was carefully monitored to avoid erroneous and over-fitting of spectra.

**Figure 4:**
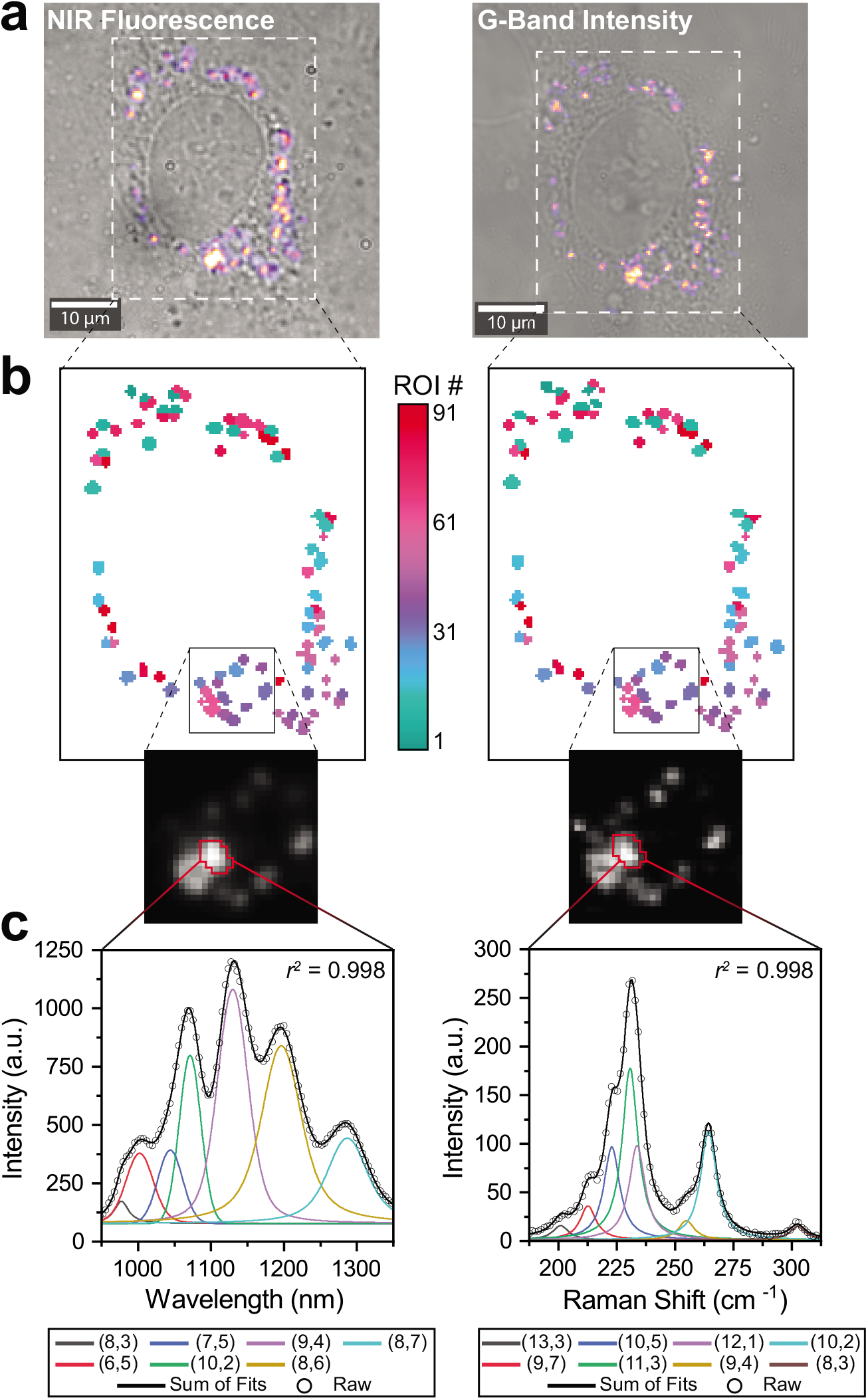
Colocalization of single-cell NIR fluorescence and Raman signals. **(a)** Transmitted light and brightfield images merged with broadband fluorescence and G-band intensity images, respectively, of a single cell incubated with DNA-SWCNTs. **(b)** Segmented ROI mask images determined from the fluorescence and G-band intensity images in (a). Inset shows magnified fluorescence and G-band intensity pixels corresponding to the indicated region in the masked image. **(c)** Deconvoluted fluorescence spectrum and RBM range of the Raman spectrum from the outlined cellular ROI in (b). Peaks from the fluorescence and RBM spectrum were fit to Voigt or Lorentz line shapes, respectively.

### Fluorescence Modulation of Concentration Subcellular Regions

With the highly improved spatial resolution, we first revisited the correlation between fluorescence intensity and G-band intensity using the colocalized subcellular ROIs. Figure 5a shows fluorescence intensity as a function of G-band intensity of all intracellular ROIs containing (GT)_30_-SWCNTs at various time points. A linear regression was performed, and the Pearson coefficient (*r_p_*) was calculated for each dataset, revealing a statistically significant positive correlation (*r_p_* > 0.4, *p* < 1e-4) at each time point. The sustained positive relationship between fluorescence intensity and DNA-SWCNT concentration confirmed a common mechanism could explain their temporal fluctuations, including the simultaneous increases observed from 0h to 6h. Since it is not possible for more DNA-SWCNTs to enter the cells after their initial dosing, we conclude that this is a signal of vesicle coalescence due to progression of the endolysosomal pathway.

We next investigated the relationship between sub-cellular DNA-SWCNT concentration and fluorescence emission modulation. Hyperspectral maps of (9,4)-SWCNT emission wavelength and G-band intensity were constructed from the ROIs of cells imaged at 0h or 6h (Fig. 5b,c), revealing that red shifted regions were generally correlated to more concentrated areas regardless of incubation time. To quantify this trend, scatter plots were created to compare these two measurements from all ROIs at 0h or 6h time points (Fig. 5d,e). Average values from 0h data were used to split the ROI population into four quadrants, thus providing a quantitative measure of their change due to intracellular processing events. The majority of DNA-SWCNT-containing ROIs had simultaneously red shifted and increased in concentration after 6h, however these shifts were completely prevented in cells treated with NOC or CQ (Fig. S7), confirming the mechanism related to vesicle trafficking processes. The same trend was observed comparing the G-band intensity to (8,6)-SWCNT emission wavelength (Fig. S9), revealing no apparent dependencies on SWCNT chirality. We partially attribute this correlation to the formation of DNA-SWCNT-protein aggregates once the nanotube concentration reaches a certain threshold, in which densely packed proteins can increasingly perturb the DNA wrapping to increase accessibility of the nanotube surface. This ultimately modulates the local dielectric environment, thus red shifting the SWCNT emission.^35, 57^ At the same time, mature late endosomes and endolysosomes undergo a series of changes in their luminal environments, including a fluctuation of ion concentrations and an increase in negative surface charges,^3^ which could further decrease the fluorescence emission energy.^36, 38^ We surmise that a combination of these factors could be contributing to the observed ROI characteristics, however the dramatic transition of DNA-SWCNT optical properties, coupled with previous reports of endolysosome localization over 6h,^5, 19^ strongly suggests that these features could be the result of specific vesicle maturation stages within lysosomal organelles.

**Figure 5:**
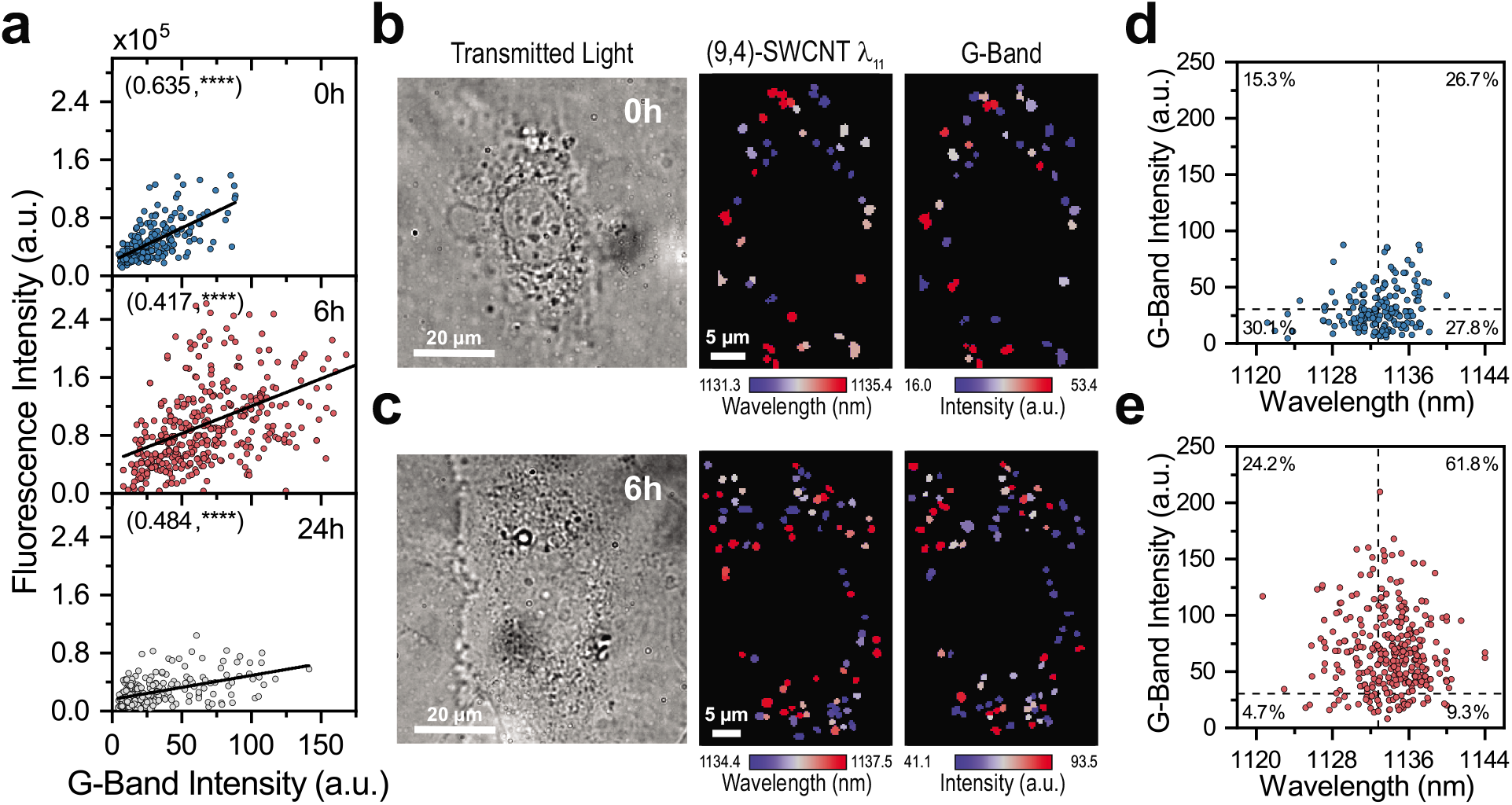
Fluorescence modulation from DNA-SWCNTs within concentrated subcellular regions. **(a)** (GT)_30_-SWCNT fluorescence intensity as a function of G-band intensity from all intracellular ROIs, with linear fits, at indicated time points. Pearson correlation coefficients, displayed in parentheses, were calculated from scatter data at each time point. Transmitted light images, (9,4)-SWCNT emission maps, and G-band intensity maps of individual cells at **(b)** 0h or **(c)** 6h time points. Color scale range encompasses 20 – 80% of values from each ROI map. **(d)** G-band intensity as a function of (9,4)-SWCNT emission wavelength from all 0h or **(e)** 6h intracellular ROIs. Average values from 0h data, represented as dashed lines, were used to compute the percent of ROIs in each quadrant.

### Intracellular Aggregate Formation is Time Dependent

The RBM of a DNA-SWCNT Raman spectrum is directly related to the resonance of a chirality’s transition energy with the excitation laser source.^28^ In the case of SWCNT aggregation, a global decrease of transition energies causes distinctive changes to components of the RBM spectrum (Fig. S11).^31^ More specifically, we observed that chiralities with *E_22_* < *E_laser_* (*E_22_* > *E_laser_*) displayed a substantial intensity decrease (increase) upon aggregation in control experiments (Fig. S12a), providing a viable basis to probe the dynamics of intracellular aggregation using fitted RBM data (Fig. 6a). To visualize the temporal progression of chirality components in all ROI spectra, we constructed a heat map illustrating the relative intracellular intensity change of each chirality with respect to solution intensities (Fig. 6b) and included the aggregated controls as a reference. Each chirality was grouped based on its solution *E_22_* value, revealing a clear trend as almost every chirality experienced an intensity change that suggested some amount of intracellular aggregation.

**Figure 6:**
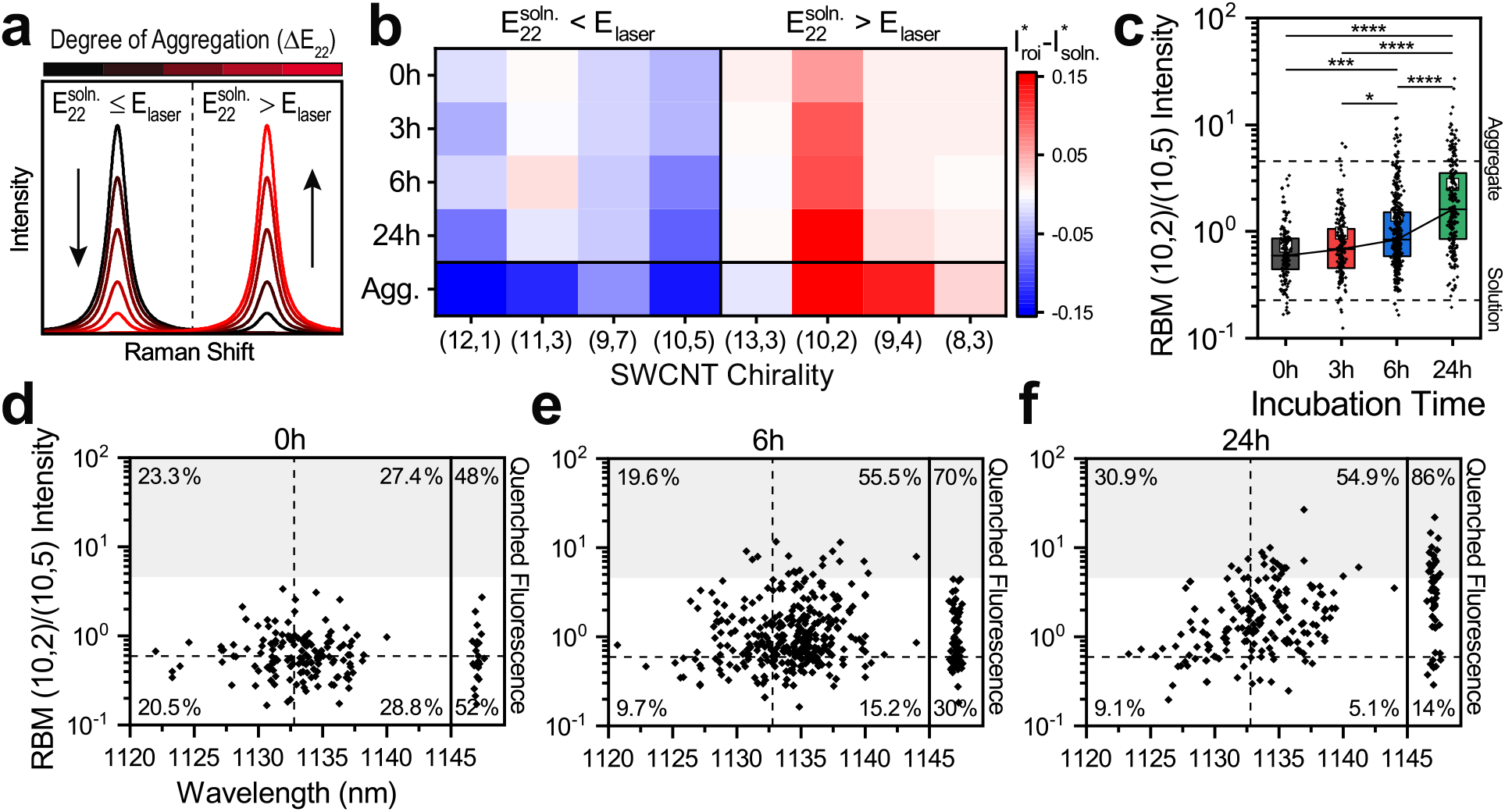
Intracellular aggregate formation is time dependent. **(a)** The RBM peak intensity of a single SWCNT depends on its transition energy (*E_22_*) and the excitation energy (*E_laser_*). Aggregation shifts the optical transition to lower energies (Δ *E_22_*), resulting in selective intensity enhancement for chiralities brought into resonance (*E_22_^soln^* ≥ *E_laser_*) and intensity reduction for chiralities brought out of resonance (*E_22_^soln^* ≥ *E_laser_*) with the excitation. **(b)** Heat map representing the change of intracellular (GT)_30_-SWCNT RBM intensities from solution as a function of chirality and time. Control intensities of intentionally aggregated (GT)_30_-SWCNTs are displayed as a reference. The chirality intensities from each ROI or control replicate were normalized by the total RBM intensity and average values are reported. **(c)** The ratio of RBM (10,2)/(10,5) intensities of all intracellular ROIs as a function of time. Boxes represent 25-75% of the data, small white squares represent the mean, trend lines connect medians, and dashed lines indicate values from aggregated or solution controls. One-way ANOVA with Tukey post hoc analysis was performed (\**p* < 0.05, \*\**p* < 0.01, \*\*\**p* < 0.001, ****p < 1e-4). The ratio of RBM (10,2)/(10,5) intensities as a function of (9,4)-SWCNT emission wavelength of all **(d)** 0h, **(e)** 6h, or **(f)** 24h ROIs. Boxed column scatter plots on the right-hand side depict RBM ratio values from ROIs with poorly fitting or quenched fluorescence. Median values from 0h data, represented as dashed lines, were used to compute the percent of ROIs in each quadrant. Shaded regions indicate the RBM (10,2)/(10,5) intensity threshold identified from aggregated controls.

Next, we devised an intracellular aggregation measurement based on the RBM intensity changes observed upon SWCNT aggregation. We identified the ratiometric RBM intensity of (10,2)/(10,5) as a suitable metric for a number of reasons: (1) distinguishable RBM peaks from both chiralities are present in aggregated and solution controls, (2) the *E_22_* of (10,5)-SWCNTs in solution (~1.58 eV) is directly in resonance with the laser and can only decrease with aggregate formation, (3) the *E_22_* of (10,2)-SWCNTs in solution (~1.69 eV) is greater than the laser energy and would move into resonance upon aggregate formation, however the expected shift (~70 meV)^58^ due to complete bundling could not decrease the transition below the laser energy. Therefore, the RBM (10,2)/(10,5) intensity ratio (hereby referred to as the “RBM aggregate ratio”) could directly relate chirality intensities to their transition energies to provide a quantitative measure of aggregation. Significantly different values of the RBM aggregate ratio were calculated from DNA-SWCNT controls of solution and aggregated spectra (Fig. S12b), providing reference points to compare against the cellular data. The RMB aggregate ratio was then calculated for every intracellular ROI containing (GT)_30_-SWCNTs and box plots were constructed for the full dataset (Fig. 6c), revealing significant differences between the distributions which increased and broadened in time. We then investigated whether a relationship could be identified between the degree of aggregation, environmental conditions within ROIs, and time of DNA-SWCNT processing within the cells. Scatter plots of the RBM aggregate ratio as a function of (9,4)-SWCNT emission wavelength were constructed from ROIs after 0h, 6h, or 24h of incubation with internalized (GT)_30_-SWCNTs (Fig. 6d-f). Again, each set of ROIs were split into four populations based on median values from 0h data. The percentage of ROIs with red shifted and increased RBM ratios effectively doubled from 0h to 6h, yet this number plateaued with additional incubation time. At the same time, an increasing number of ROIs became quenched over time (Fig. S13). The majority of quenched ROIs, however, exhibited substantial aggregation, as shown in the righthand column scatter plots of each time point. These spectral characteristics could be indicative of increasingly harsh environmental conditions which the DNA-SWCNTs were subjected to during later stages in the processing pathway, as evidenced by sequential red shifting, aggregate formation, and fluorescence quenching due to excessive aggregation.^49, 50^ Furthermore, these changing dynamics establish a potential end point to the intracellular processing pathway, at which the extended time within the degradative lysosomal environment results in irreversible bundling of internalized DNA-SWCNTs.

### Multispectral Fingerprinting of Vesicle Trafficking

Vesicle trafficking is naturally an asynchronous process, and thus a group of ROIs collected from cells at a specific time is characterized by heterogeneity due to the continuum of processes occurring at a given moment. Therefore, we sought to characterize the full ROI dataset simultaneously using a spectral *k*-means clustering algorithm,^59^ which would identify four groups (clusters) that commonly appeared regardless of incubation time. First, the normalized RBM spectrum from all intracellular ROIs were pooled together from cells dosed with (GT)_30_-SWCNTs and incubated for 0h, 3h, 6h, or 24h, and input to a spectral *k*-means clustering code (Fig. 7a). Briefly, the matrix of RBM spectra was projected to a lower-dimensional representation through a series of mathematical transformations, allowing for key spectral components to be identified while exaggerating the differences between the set of ROIs. Then, the transformed data was segmented using a standard *k*-means clustering analysis,^60^ resulting in four ROI populations which could best represent the full dataset. The average RBM spectrum (Fig. 7b) and fluorescence spectrum (Fig. 7c) from ROIs assigned to each cluster were distinct, enabling four intracellular spectral fingerprints to be identified based on the properties of ROIs in each cluster. Box plots were constructed to identify and compare spectral characteristics. The DNA-SWCNT concentration (Fig. 7d) and fluorescence intensity (Fig. 7e) both fluctuated up and down from cluster 1 to 4 in a similar trend. In contrast, intracellular aggregation (Fig. 7f) sharply increased from cluster 2 to 4 while environmental changes within the ROIs (Fig. 7f), determined by emission center wavelengths of the (9,4)-SWCNTs, monotonically increased from cluster 1 to 4. Although these clusters were determined independent of DNA-SWCNT incubation time, we identified a temporal association between the percentage of ROIs in each cluster and increasing length of intracellular processing (Fig. 7h). Clusters 1 and 2 were more prevalent in earlier time points while clusters 3 and 4 monotonically increased with increasing incubation time, suggesting a relationship between cell cluster composition and time of intracellular processing.

**Figure 7:**
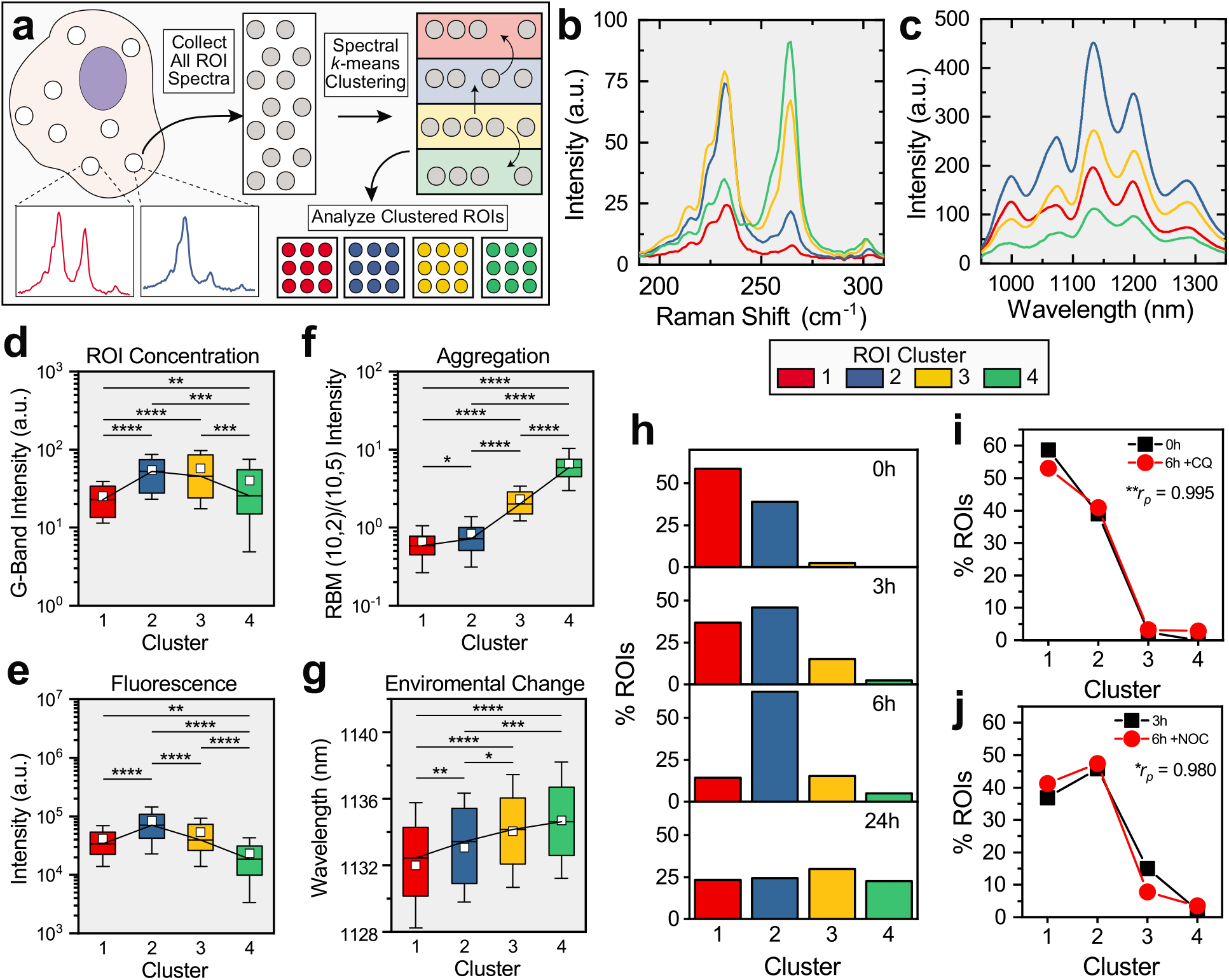
Spectral *k*-means clustering facilitates vesicle mapping *via* DNA-SWCNT spectral fingerprint. **(a)** Schematic depicting the process of *k*-means clustering. The normalized RBM spectra of (GT)_30_-SWCNTs in all cellular ROIs were pooled together across time points and input to a spectral *k*-means clustering algorithm. Each ROI was assigned to one of four clusters based on population similarities, and the ROIs identified to each cluster were collected and analyzed independently (see methods for full details). **(b)** The average RBM spectrum and **(c)** fluorescence spectrum of each computed cluster. Box plots depicting **(d)** G-band intensity, **(e)** fluorescence intensity, **(f)** RBM (10,2)/(10,5) intensity, and **(g)** (9,4)-SWCNT emission wavelength of ROIs classified to each cluster. Boxes represent 25-75% of the data, small white squares represent means, trend lines connect medians, and whiskers represent mean ± s.d. One-way ANOVA with Tukey post hoc analysis was performed for each metric**. (h)** The percent of ROIs at each time point that were classified into each cluster. **(i)** The percent of ROIs assigned to each cluster from cells treated with CQ or **(j)** NOC for 6h. ROIs were matched to one of the four clusters defined in (b) and the Pearson coefficient (*r_p_*) was calculated to quantify the correlation between treated cells to untreated cells at indicated time points. (\**p* < 0.05, \*\**p* < 0.01, \*\*\**p* < 0.001, \*\*\*\**p* < 1e-4 for all statistical measurements).

To further probe the dynamics related to vesicle maturation, we segmented the ROIs from pharmacological inhibitor-treated cells into the four pre-defined populations using a *k*-nearest neighbor distance query,^61^ which identified each ROI’s closest matching cluster based on the distance between points of their normalized RBM spectrum. The resulting cluster percentages of CQ and NOC treated cells (Fig. 7i,j) were nearly identical to the untreated cells from 0h or 3h, respectively, further confirming the role of vesicle maturation in the spectral evolution of internalized DNA-SWCNTs. Therefore, we believe that each cluster can be interpreted as a sequential step in the intracellular trafficking pathway, in which progression of vesicle maturation can be determined based on the spectral fingerprint of encapsulated DNA-SWCNTs.

To illustrate these findings in terms of sequential trafficking events, we propose a schematic (Fig. 8) to describe the intracellular processes which DNA-SWCNTs are subjected to over the course of 24h. The DNA-SWCNTs first enter the cell through active endocytic processes (i), where they are retained within membrane-derived vesicles for the duration of their subsequent pathway. Fusion of DNA-SWCNT-containing vesicles (ii) drives a concentration increase in discrete localized regions, where accumulated DNA-SWCNTs exhibit EET as they become tightly packed together. At the same time, the vesicle maturation process is initiated to produce progressively degradative conditions (iii). This process causes conformational changes to the DNA wrapping, promotes increasingly complex interactions between the DNA-SWCNTs and luminal biomolecules, and ultimately modulates the dielectric environment to induce a red shift of DNA-SWCNT fluorescence emission. After an extended period of time in the harsh environment (iv), internalized DNA-SWCNTs aggregate within endolysosomal organelles and fluorescence emission is quenched, marking the end of the trafficking pathway. In contrast, pharmacological inhibition of vesicle maturation effectively suppresses these key spectral changes by obstructing initial vesicle fusion and preventing (CQ) or limiting (NOC) environmental transformations.

**Figure 8:**
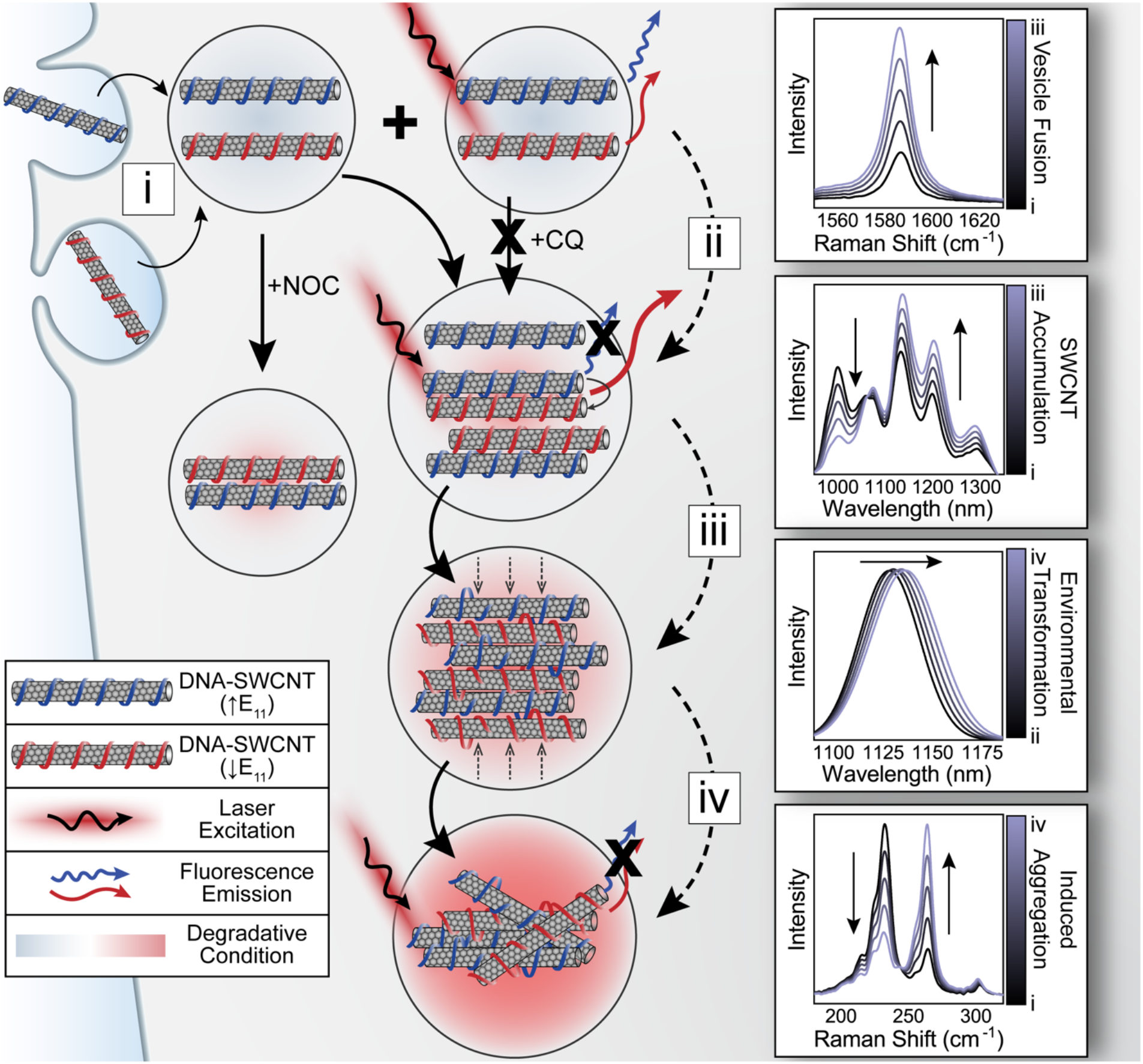
Schematic of the spectral evolution which characterizes DNA-SWCNT trafficking. DNA-SWCNTs are internalized by HUVEC cells *via* active endocytic processes (i), where they are contained in vesicles derived from the plasma membrane. Vesicles containing small amounts of DNA-SWCNTs eventually fuse (ii) to drive an increase in the local concentration and cause exciton energy transfer (EET) between closely packed DNA-SWCNTs. Vesicle fusion can be inhibited using Chloroquine (CQ) or Nocodazole (NOC), however NOC does not stop initial vesicle acidification. Luminal conditions become increasingly harsh (iii) as vesicles mature, inducing red-shifted fluorescence emission as the DNA wrapping is increasingly perturbed, and eventually causing hard-contact DNA-SWCNT aggregation (iv).

## Conclusions

The results obtained from the spectral fingerprinting method provide a unique perspective on the intracellular processing pathways which internalized DNA-SWCNTs traverse. The fluorescence and Raman spectra from whole cells were first examined, revealing an increase of fluorescence and G-band intensities from 0h to 6h of equal magnitude. At the same time, intensity changes from fluorescence emission bands suggested the occurrence of EET between closely packed DNA-SWCNT chiralities. Although the extent of EET plateaued at 6h, direct aggregation of internalized DNA-SWCNTs was indicated by changes in RBM band intensities, which monotonically scaled with incubation time. To confirm these events were induced by progression of the intracellular trafficking pathway, cells were treated with two pharmacological inhibitors of vesicle maturation, both of which suppressed the identified spectral changes over 6h *via* distinct mechanisms of action.

To investigate these processes at the subcellular level, we developed a segmentation method which could colocalize the Raman and fluorescence spectrum of internalized DNA-SWCNTs within nanoscale regions. A simultaneous multi-peak fitting algorithm, which provided single-chirality resolution of multicomponent spectra, was used to quantify the relevant spectral features and characterize the conditions of each cellular ROI. This approach determined correlations between fluorescence intensity, DNA-SWCNT concentration, fluorescence emission wavelength, and aggregate formation within cellular ROIs for the first time, illustrating the effect of changing intracellular conditions on the internalized DNA-SWCNTs. Finally, we developed a method to identify various stages of the vesicle trafficking process based on the responsive optical properties of internalized DNA-SWCNTs. A spectral *k*-means clustering algorithm was employed to generate four spectral fingerprints which could describe both the physical state of the DNA-

SWCNTs and the environmental conditions within a given ROI. The temporal fluctuation of each population revealed their associations with specific steps in the intracellular processing pathway, suggesting the progress of a single vesicle could be estimated based on the spectral fingerprint of encapsulated DNA-SWCNTs. Furthermore, the temporal evolution of each spectral component delineated the significant steps in the vesicle maturation process, enabling the comprehensive characterization of DNA-SWCNT progression though intracellular trafficking pathways. The importance of ENM surface chemistry on their biological interactions implicates these findings in a myriad of ENM systems, including those which utilize a biomolecular surface functionalization. Moreover, the data processing pipeline detailed here could be adapted for any optically active SWCNT formulation, providing a fully realized approach to enable specific SWCNT properties to be investigated within complex biological environments.

## Materials and Methods

### DNA-SWCNT Sample Preparation

Raw single-walled carbon nanotubes produced by the HiPco process (Nanointegris) were used throughout this study. For each dispersion, 2 mg of (GT)_6_ or (GT)_30_ oligonucleotide (Integrated DNA Technologies) were added to 1 mg of raw nanotubes, suspended in 1 mL of 0.1M NaCl (Sigma-Aldrich), and ultrasonicated using a 1/8” tapered microtip for 30 min at 40% amplitude (Sonics Vibracell VCX-130; Sonics and Materials). The resultant suspensions were ultracentrifuged (Sorvall Discovery M120 SE) for 30 min at 250,000 × g and the top ~80% of the supernatant was collected. Concentrations were determined using a UV/vis/NIR spectrophotometer (Jasco, Tokyo, Japan) and the extinction coefficient of A910 = 0.02554 L mg^−1^ cm^−1^.^55^

### Cell Culture

HUVEC cells (ATCC, Manassas, VA, USA) were cultured under standard incubation conditions at 37 °C and 5% CO2 in endothelial growth media (EGM BulletKit CC-3124, Lonza). For all imaging experiments, cells were seeded into grid labeled collagen-coated 35mm glass bottom microwell dishes (MatTek) to a final concentration of 5,000 cells/ cm^2^ and allowed to culture for at least 48 hours, with regular media replacement every 24 hours. To dose the cells, the media was removed from each culture dish, replaced with 1 mg-L^−1^ (GT)_6_-SWCNT or (GT)_30_-SWCNT diluted in media,^20^ and incubated for 1 hour to allow internalization into the cells. The SWCNT-containing media was removed, the cells were rinsed 3X with sterile phosphate buffered saline (PBS, Gibco), and fresh media was replenished. The 0h samples were immediately fixed using 4% paraformaldehyde in PBS for 10 minutes, rinsed 3X with PBS, and covered with PBS to retain an aqueous environment during imaging. The 3h, 6h, and 24h samples were later fixed using the same procedure.

### Near-Infrared Fluorescence Microscopy

A near-infrared hyperspectral fluorescence microscope, similar to a previously described system,^55^ was used to obtain the hyperspectral fluorescence images from fixed cell samples. Briefly, a continuous 730 nm diode laser with 1.5 W output power was injected into a multimode fiber to produce an excitation source, which was reflected on the sample stage of an Olympus IX-73 inverted microscope equipped with a UApo N 100× /1.49 oil immersion IR objective (Olympus, USA). Emission was passed through a volume Bragg Grating and collected with a 2D InGaAs array detector (Photon Etc.) to generate spectral image stacks. Fixed cell samples were mounted on the hyperspectral microscope to obtain transmitted light images and hyperspectral images from internalized DNA-SWCNTs in individual cells at each time point. Hyperspectral data were processed and extracted using custom codes written with Matlab software.

### Confocal Raman Microscopy

Each cell sample was imaged with an inverted WiTec Alpha300 R confocal-Raman microscope (WiTec, Germany) equipped with a Zeiss EC Epiplan-Neofluar 100× /0.9 oil objective, a 785 nm laser source set to 35 mW sample power, and collected with a UHTS 300 spectrograph (600 lines/mm grating) coupled with an Andor DR32400 CCD detector (−61 °C, 1650 x 200 pixels). Small cellular areas were scanned, and spectra were obtained in 0.29 × 0.29 μm intervals using 0.2 s integration time per spectrum to construct hyperspectral images of individual cells. Global background subtraction and cosmic-ray removal were performed on each scan using Witec Project 5.2 software. Hyperspectral data was extracted and processed using custom codes written with Matlab software.

### Pharmacological Inhibition of Endosomal Maturation

HUVEC cells were cultured and dosed with (GT)_6_-SWCNTs or (GT)_30_-SWCNTs following the same procedure previously described, however the media used to replenish the cells after DNA-SWCNT removal and PBS rinsing was spiked with 10 μg-mL^−1^ Nocodazole (NOC) or 100 μM Chloroquine (CQ). The cells were incubated for 6 hours following the addition of inhibitors before fixation in 4% paraformaldehyde in PBS for 10 minutes. The cells were then imaged following the same procedure used for the untreated cells.

### ROI Colocalization

ROI colocalization was carried out on all hyperspectral ‘cubes’ (*i.e*., three-dimensional datasets in which x and y dimensions are spatial coordinates, the z dimension is the spectral coordinate, and the pixel value corresponds to the spectral intensity) following initial background subtraction and cosmic-ray removal steps. Using custom Matlab codes, composite fluorescence images were created by integrating the entire spectral dimension and composite Raman images were created by integrating the G-band or RBM regions of the spectrum. The fluorescence and Raman images were first roughly colocalized by applying an intensity threshold to each image, binarizing and segmenting each image individually, determining the intensity-weighted centroid of each segmented ROI, and iteratively overlaying the images to find the coordinates which minimize the root-mean-square deviation (RMSD) of similar ROIs. Next, the composite images and cubes were cropped and imported to the open-source image processing software FIJI. The G-band composite images were segmented into ROIs, which were manually adjusted to ensure consistency in the segmentation process, and compared with the RBM region of the Raman cube. The ROIs determined from the Raman data were then transferred to the fluorescence images, where each ROI was manually adjusted to account for minor discrepancies in their location and shape. Once all ROIs were determined for a cell, their locations were imported to Matlab for further analysis. Of note, the resolution of the confocal Raman area scans was experimentally optimized prior to data acquisition to match the pixel size of the hyperspectral fluorescence microscope, and thus the spatial resolution of the two cubes was essentially the same. ROI location adjustments mainly accounted for minor rotations of the imaging field as the result of mounting on two separate instruments, and ROIs which could not be clearly identified as the same were disregarded. In certain cases, ROIs which exhibited strong Raman intensities in both the G-band and RBM regions displayed little to no fluorescence intensity from the same spatial location. These ROIs were considered to be colocalized and accurate only if their G-band and RBM integrated intensities were comparable with other ROIs in the same image and other nearby ROIs which exhibited fluorescence were colocalized with Raman signal. The appearance of visible, dark spots within these ROIs in transmitted light images obtained with the hyperspectral fluorescence microscope were also used to verify the presence of DNA-SWCNTs which were quenched. An example of quenched fluorescence from cellular ROIs is provided in the supporting information. Any and all ROIs which could not be definitively colocalized were disregarded from further data analysis.

### Multi-Peak Fitting of the Fluorescence and RBM Spectra

The colocalized ROI data for the fluorescence and Raman cubes were used to obtain average spectra from each ROI, which was processed with a custom Matlab pipeline. First, the average fluorescence and Raman spectrum from each ROI were calculated by averaging pixel intensity values in their x-y direction and extracting the spectral z dimension from each cube. The fluorescence spectrum from each ROI was fitted to an additive combination of Voigt line shapes corresponding to the single chirality component spectra, and only chiralities which were identified to significantly contribute to the fluorescence spectrum were included in the fitting process. The peak center wavelength and width parameters of each chirality were allowed to vary independently, but each parameter was limited within the same set of constraints. The area under the curve and global offset were restricted to non-negative values. The radial breathing mode of the Raman spectrum in each ROI was fit to an additive combination of Lorentz line shapes corresponding to the single chirality component spectra. The chiralities which were included in the fits were chosen based on (1) their resonance with the excitation laser, determined by empirical Katura plots found in the literature,^48^ (2) their presence when spectra were obtained from solution controls, and (3) their presence when spectra were obtained from aggregated samples. Peak centers were initially specified and allowed to shift within a very small window, however each spectrum was restricted to a single full width and half maximum for all peaks.^56^ The area under the curve and global offset were restricted to non-negative values. *r^2^* > 0.95 was used as a cutoff to remove poor-fitting ROI data from further analyses.

### Spectral k-means Clustering

Spectral *k*-means clustering of intracellular ROIs was performed in Matlab using a combination of custom and toolbox functions. Clustering was first carried out using ROI data from cells dosed with (GT)_30_-SWCNTs and incubated for 0h, 3h, 6h, or 24h. The RBM region of the average Raman spectrum in each ROI was collected, normalized by the total RBM intensity, and compiled into a matrix containing the full dataset. This matrix was used as an input to a spectral clustering algorithm, which mathematically identified and connected groups of similar datapoints using a well-defined multi-step process.^59, 62^ Briefly, the matrix of ROI spectra was transformed to a lower dimension by calculating the Euclidian distance similarity matrix, determining the normalized random-walk Laplacian matrix, and solving for the *k* smallest eigenvalues to construct a new matrix with *k* datapoints to represent the spectrum from each ROI. The resulting matrix was segmented into 4 clusters using a standard *k*-means clustering algorithm,^60^ and the full set of data corresponding to each ROI was collected and grouped by cluster identification for further analysis. Existing clusters were appended with new ROI data using a *k*-nearest neighbor distance query, in which the normalized RBM spectrum from each new ROI was compared to the average normalized RBM spectrum from all four clusters, Euclidian distances were calculated for each, and the ROI was assigned to the nearest neighbor (minimized distance) cluster.

### Statistical Analysis

OriginPro 2018 was used to perform all statistical analysis. All data either met assumptions of the statistical tests performed (*i.e*., normality, equal variances, etc.) or was transformed to meet assumptions before statistical analysis was carried out. Statistical significance was analyzed using two-sample two-tailed student t-test or one-way ANOVA where appropriate. Testing of multiple hypotheses was accounted for by performing one-way ANOVA with Tukey’s post hoc test. Specific information about statistical analyses can be found in figure legends.

## Acknowledgements

This work was supported by National Science Foundation CAREER Award # 1844536, the RI-INBRE Early Career Development Award Grant # P20GM103430 from the National Institute of General Medical Sciences of the National Institutes of Health, the Rhode Island Foundation – Medical Research Fund, and the URI College of Engineering. The confocal Raman data was acquired at the RI Consortium for Nanoscience and Nanotechnology, a URI College of Engineering core facility partially funded by the National Science Foundation EPSCoR, Cooperative Agreement #OIA-1655221. We acknowledge and thank P. V. Jena for the Matlab codes which were adapted to become a part of the full data processing pipeline.

## Supporting Information

Figure S1. Solution-based optical characterization of DNA-SWCNTs.

Figure S2. Fluorescence intensity and local concentration of (GT)_6_-SWCNTs are co-dependent within single cells.

Figure S3. RBM of DNA-SWCNTs in solution and aggregated.

Figure S4. Temporal resolution of (GT)_6_-SWCNT spectral features indicates aggregation within subcellular regions.

Figure S5. Spectral response to inhibition of endosomal progression.

Figure S6. Fluorescence modulation from (GT)_6_-SWCNTs within concentrated subcellular regions.

Figure S7. G-band intensity of (GT)_30_-SWCNTs as a function of (9,4)-SWCNT emission wavelength in inhibitor-treated cells.

Figure S8. G-band intensity of (GT)_6_-SWCNTs as a function of (9,4)-SWCNT emission wavelength in inhibitor-treated cells.

Figure S9. G-band intensity of (GT)_30_-SWCNTs as a function of (8,6)-SWCNT emission wavelength.

Figure S10. G-band intensity of (GT)_6_-SWCNTs as a function of (8,6)-SWCNT emission wavelength.

Figure S11. Spectral deconvolution of DNA-SWCNTs in solution and aggregated.

Figure S12. RBM chirality intensities from aggregated DNA-SWCNTs.

Figure S13. Quenched fluorescence of DNA-SWCNTs in cellular ROIs.

Figure S14. Intracellular aggregate formation is time dependent.

Figure S15. *k*-means clustering assignment of (GT)_6_-SWCNTs

Table S1: Estimate of the optical properties of DNA-SWCNT chiralities identified in the fluorescence spectrum when excited by a 730 nm laser.

Table S2: Estimate of the optical properties of DNA-SWCNT chiralities identified in the RBM region of the Raman spectrum when excited by a 1.58 eV laser.

## Notes

### Competing Interest Statement

The authors have declared no competing interest.

## References

1. Conner, S.D.; Schmid, S. L., Regulated portals of entry into the cell. Nature 2003, 422 (6927), 37–44.

2. Rink, J.; Ghigo, E.; Kalaidzidis, Y.; Zerial, M., Rab conversion as a mechanism of progression from early to late endosomes. Cell 2005, 122 (5), 735–749.

3. Huotari, J.; Helenius, A., Endosome maturation. The EMBO journal 2011, 30 (17), 3481–3500.

4. Oh, N.; Park, J.-H., Endocytosis and exocytosis of nanoparticles in mammalian cells. International journal of nanomedicine 2014, 9 (Suppl 1), 51.

5. Jena, P. V.; Roxbury, D.; Galassi, T. V.; Akkari, L.; Horoszko, C. P.; Iaea, D. B.; Budhathoki-Uprety, J.; Pipalia, N.; Haka, A. S.; Harvey, J. D., A carbon nanotube optical reporter maps endolysosomal lipid flux. ACS nano 2017, 11 (11), 10689–10703.

6. Xu, H.; Ren, D., Lysosomal physiology. Annual review of physiology 2015, 77, 57–80.

7. Donahue, N. D.; Acar, H.; Wilhelm, S., Concepts of nanoparticle cellular uptake, intracellular trafficking, and kinetics in nanomedicine. Advanced drug delivery reviews 2019, 143, 68–96.

8. Chithrani, B. D.; Ghazani, A. A.; Chan, W. C., Determining the size and shape dependence of gold nanoparticle uptake into mammalian cells. Nano letters 2006, 6 (4), 662–668.

9. Dasgupta, S.; Auth, T.; Gompper, G., Shape and orientation matter for the cellular uptake of nonspherical particles. Nano letters 2014, 14 (2), 687–693.

10. Zhu, M.; Nie, G.; Meng, H.; Xia, T.; Nel, A.; Zhao, Y., Physicochemical properties determine nanomaterial cellular uptake, transport, and fate. Accounts of chemical research 2013, 46 (3), 622–631.

11. Walkey, C. D.; Olsen, J. B.; Guo, H.; Emili, A.; Chan, W. C., Nanoparticle size and surface chemistry determine serum protein adsorption and macrophage uptake. Journal of the American Chemical Society 2012, 134 (4), 2139–2147.

12. Pinals, R. L.; Yang, D.; Rosenberg, D. J.; Chaudhary, T.; Crothers, A. R.; Iavarone, A. T.; Hammel, M.; Landry, M. P., Quantitative Protein Corona Composition and Dynamics on Carbon Nanotubes in Biological Environments. Angewandte Chemie 2020.

13. Nixon, R. A., Endosome function and dysfunction in Alzheimer’s disease and other neurodegenerative diseases. Neurobiology of aging 2005, 26 (3), 373–382.

14. Fukuda, T.; Ewan, L.; Bauer, M.; Mattaliano, R. J.; Zaal, K.; Ralston, E.; Plotz, P. H.; Raben, N., Dysfunction of endocytic and autophagic pathways in a lysosomal storage disease. Annals of Neurology: Official Journal of the American Neurological Association and the Child Neurology Society 2006, 59 (4), 700–708.

15. Sarkar, S.; Carroll, B.; Buganim, Y.; Maetzel, D.; Ng, A. H.; Cassady, J. P.; Cohen, M. A.; Chakraborty, S.; Wang, H.; Spooner, E., Impaired autophagy in the lipid-storage disorder Niemann-Pick type C1 disease. Cell reports 2013, 5 (5), 1302–1315.

16. Mercer, J.; Schelhaas, M.; Helenius, A., Virus entry by endocytosis. Annual review of biochemistry 2010, 79, 803–833.

17. Sandin, P.; Fitzpatrick, L. W.; Simpson, J. C.; Dawson, K. A., High-speed imaging of Rab family small GTPases reveals rare events in nanoparticle trafficking in living cells. ACS nano 2012, 6 (2), 1513–1521.

18. Vercauteren, D.; Deschout, H.; Remaut, K.; Engbersen, J. F.; Jones, A. T.; Demeester, J.; De Smedt, S. C.; Braeckmans, K., Dynamic colocalization microscopy to characterize intracellular trafficking of nanomedicines. ACS nano 2011, 5 (10), 7874–7884.

19. Galassi, T. V.; Jena, P. V.; Shah, J.; Ao, G.; Molitor, E.; Bram, Y.; Frankel, A.; Park, J.; Jessurun, J.; Ory, D. S., An optical nanoreporter of endolysosomal lipid accumulation reveals enduring effects of diet on hepatic macrophages in vivo. Science translational medicine 2018, 10 (461), eaar2680.

20. Gravely, M.; Safaee, M. M.; Roxbury, D., Biomolecular Functionalization of a Nanomaterial To Control Stability and Retention within Live Cells. Nano letters 2019, 19 (9), 6203–6212.

21. Jin, S.; Wijesekara, P.; Boyer, P. D.; Dahl, K. N.; Islam, M. F., Length-dependent intracellular bundling of single-walled carbon nanotubes influences retention. Journal of Materials Chemistry B 2017, 5 (32), 6657–6665.

22. Kneipp, J.; Kneipp, H.; Wittig, B.; Kneipp, K., Following the dynamics of pH in endosomes of live cells with SERS nanosensors. The Journal of Physical Chemistry C 2010, 114 (16), 7421–7426.

23. Huefner, A.; Kuan, W.-L.; Müller, K. H.; Skepper, J. N.; Barker, R. A.; Mahajan, S., Characterization and visualization of vesicles in the endo-lysosomal pathway with surface-enhanced Raman spectroscopy and chemometrics. ACS nano 2016, 10 (1), 307–316.

24. Gowen, A. A.; O’Donnell, C. P.; Cullen, P. J.; Downey, G.; Frias, J. M., Hyperspectral imaging–an emerging process analytical tool for food quality and safety control. Trends in food science & technology 2007, 18 (12), 590–598.

25. Jorio, A.; Fantini, C.; Pimenta, M.; Heller, D.; Strano, M.; Dresselhaus, M.; Oyama, Y.; Jiang, J.; Saito, R., Carbon nanotube population analysis from Raman and photoluminescence intensities. Applied Physics Letters 2006, 88 (2), 023109.

26. Ebbesen, T.; Lezec, H.; Hiura, H.; Bennett, J.; Ghaemi, H.; Thio, T., Electrical conductivity of individual carbon nanotubes. Nature 1996, 382 (6586), 54–56.

27. Bachilo, S. M.; Strano, M. S.; Kittrell, C.; Hauge, R. H.; Smalley, R. E.; Weisman, R. B., Structure-assigned optical spectra of single-walled carbon nanotubes. science 2002, 298 (5602), 2361–2366.

28. Jorio, A.; Pimenta, M.; Souza Filho, A.; Saito, R.; Dresselhaus, G.; Dresselhaus, M., Characterizing carbon nanotube samples with resonance Raman scattering. New Journal of Physics 2003, 5 (1), 139.

29. O’connell, M. J.; Bachilo, S. M.; Huffman, C. B.; Moore, V. C.; Strano, M. S.; Haroz, E. H.; Rialon, K. L.; Boul, P. J.; Noon, W. H.; Kittrell, C., Band gap fluorescence from individual single-walled carbon nanotubes. Science 2002, 297 (5581), 593–596.

30. Liu, Z.; Davis, C.; Cai, W.; He, L.; Chen, X.; Dai, H., Circulation and long-term fate of functionalized, biocompatible single-walled carbon nanotubes in mice probed by Raman spectroscopy. Proceedings of the National Academy of Sciences 2008, 105 (5), 1410–1415.

31. Heller, D. A.; Barone, P. W.; Swanson, J. P.; Mayrhofer, R. M.; Strano, M. S., Using Raman spectroscopy to elucidate the aggregation state of single-walled carbon nanotubes. The Journal of Physical Chemistry B 2004, 108 (22), 6905–6909.

32. Brown, S.; Jorio, A.; Dresselhaus, M.; Dresselhaus, G., Observations of the D-band feature in the Raman spectra of carbon nanotubes. Physical Review B 2001, 64 (7), 073403.

33. Smith, A. M.; Mancini, M. C.; Nie, S., Second window for in vivo imaging. Nature nanotechnology 2009, 4 (11), 710–711.

34. Choi, J. H.; Strano, M. S., Solvatochromism in single-walled carbon nanotubes. Applied Physics Letters 2007, 90 (22), 223114.

35. Heller, D. A.; Pratt, G. W.; Zhang, J.; Nair, N.; Hansborough, A. J.; Boghossian, A. A.; Reuel, N. F.; Barone, P. W.; Strano, M. S., Peptide secondary structure modulates single-walled carbon nanotube fluorescence as a chaperone sensor for nitroaromatics. Proceedings of the National Academy of Sciences 2011, 108 (21), 8544–8549.

36. Roxbury, D.; Jena, P. V.; Shamay, Y.; Horoszko, C. P.; Heller, D. A., Cell membrane proteins modulate the carbon nanotube optical bandgap via surface charge accumulation. ACS nano 2016, 10 (1), 499–506.

37. Barone, P. W.; Strano, M. S., Reversible control of carbon nanotube aggregation for a glucose affinity sensor. Angewandte Chemie 2006, 118 (48), 8318–8321.

38. Gillen, A. J.; Kupis-Rozmysłowicz, J.; Gigli, C.; Schuergers, N.; Boghossian, A. A., Xeno nucleic acid nanosensors for enhanced stability against ion-induced perturbations. The journal of physical chemistry letters 2018, 9 (15), 4336–4343.

39. Zheng, M.; Jagota, A.; Semke, E. D.; Diner, B. A.; McLean, R. S.; Lustig, S. R.; Richardson, R. E.; Tassi, N. G., DNA-assisted dispersion and separation of carbon nanotubes. Nature materials 2003, 2 (5), 338–342.

40. Gao, Z.; Varela, J. A.; Groc, L.; Lounis, B.; Cognet, L., Toward the suppression of cellular toxicity from single-walled carbon nanotubes. Biomaterials science 2016, 4 (2), 230–244.

41. Boghossian, A. A.; Zhang, J.; Barone, P. W.; Reuel, N. F.; Kim, J. H.; Heller, D. A.; Ahn, J. H.; Hilmer, A. J.; Rwei, A.; Arkalgud, J. R., Near-Infrared Fluorescent Sensors based on Single-Walled Carbon Nanotubes for Life Sciences Applications. ChemSusChem 2011, 4 (7), 848.

42. Budhathoki-Uprety, J.; Jena, P. V.; Roxbury, D.; Heller, D. A., Helical polycarbodiimide cloaking of carbon nanotubes enables inter-nanotube exciton energy transfer modulation. Journal of the American Chemical Society 2014, 136 (44), 15545–15550.

43. Safaee, M. M.; Gravely, M.; Rocchio, C.; Simmeth, M.; Roxbury, D., DNA sequence mediates apparent length distribution in single-walled carbon nanotubes. ACS applied materials & interfaces 2018, 11 (2), 2225–2233.

44. Jena, P. V.; Safaee, M. M.; Heller, D. A.; Roxbury, D., DNA–carbon nanotube complexation affinity and photoluminescence modulation are independent. ACS applied materials & interfaces 2017, 9 (25), 21397–21405.

45. Ferrari, A. C.; Robertson, J., Interpretation of Raman spectra of disordered and amorphous carbon. Physical review B 2000, 61 (20), 14095.

46. Park, H.-J.; Zhang, Y.; Georgescu, S. P.; Johnson, K. L.; Kong, D.; Galper, J. B., Human umbilical vein endothelial cells and human dermal microvascular endothelial cells offer new insights into the relationship between lipid metabolism and angiogenesis. Stem cell reviews 2006, 2 (2), 93–101.

47. Torrens, O.; Milkie, D.; Zheng, M.; Kikkawa, J., Photoluminescence from intertube carrier migration in single-walled carbon nanotube bundles. Nano letters 2006, 6 (12), 2864–2867.

48. Weisman, R. B.; Bachilo, S. M., Dependence of optical transition energies on structure for single-walled carbon nanotubes in aqueous suspension: an empirical Kataura plot. Nano letters 2003, 3 (9), 1235–1238.

49. Reich, S.; Dworzak, M.; Hoffmann, A.; Thomsen, C.; Strano, M., Excited-state carrier lifetime in single-walled carbon nanotubes. Physical Review B 2005, 71 (3), 033402.

50. Crochet, J.; Clemens, M.; Hertel, T., Quantum yield heterogeneities of aqueous single-wall carbon nanotube suspensions. Journal of the American Chemical Society 2007, 129 (26), 8058–8059.

51. Bayer, N.; Schober, D.; Prchla, E.; Murphy, R. F.; Blaas, D.; Fuchs, R., Effect of bafilomycin A1 and nocodazole on endocytic transport in HeLa cells: implications for viral uncoating and infection. Journal of virology 1998, 72 (12), 9645–9655.

52. Ohkuma, S.; Poole, B., Fluorescence probe measurement of the intralysosomal pH in living cells and the perturbation of pH by various agents. Proceedings of the National Academy of Sciences 1978, 75 (7), 3327–3331.

53. Mauthe, M.; Orhon, I.; Rocchi, C.; Zhou, X.; Luhr, M.; Hijlkema, K.-J.; Coppes, R. P.; Engedal, N.; Mari, M.; Reggiori, F., Chloroquine inhibits autophagic flux by decreasing autophagosome-lysosome fusion. Autophagy 2018, 14 (8), 1435–1455.

54. Mesaki, K.; Tanabe, K.; Obayashi, M.; Oe, N.; Takei, K., Fission of tubular endosomes triggers endosomal acidification and movement. PloS one 2011, 6 (5), e19764.

55. Roxbury, D.; Jena, P. V.; Williams, R. M.; Enyedi, B.; Niethammer, P.; Marcet, S.; Verhaegen, M.; Blais-Ouellette, S.; Heller, D. A., Hyperspectral microscopy of near-infrared fluorescence enables 17-chirality carbon nanotube imaging. Scientific reports 2015, 5 (1), 1–6.

56. Araujo, P.; Pesce, P.; Dresselhaus, M.; Sato, K.; Saito, R.; Jorio, A., Resonance Raman spectroscopy of the radial breathing modes in carbon nanotubes. Physica E: Low-dimensional Systems and Nanostructures 2010, 42 (5), 1251–1261.

57. Heller, D. A.; Jeng, E. S.; Yeung, T.-K.; Martinez, B. M.; Moll, A. E.; Gastala, J. B.; Strano, M. S., Optical detection of DNA conformational polymorphism on single-walled carbon nanotubes. Science 2006, 311 (5760), 508–511.

58. Fantini, C.; Jorio, A.; Souza, M.; Strano, M.; Dresselhaus, M.; Pimenta, M., Optical transition energies for carbon nanotubes from resonant Raman spectroscopy: Environment and temperature effects. Physical review letters 2004, 93 (14), 147406.

59. Shi, J.; Malik, J., Normalized cuts and image segmentation. IEEE Transactions on pattern analysis and machine intelligence 2000, 22 (8), 888–905.

60. Seber, G. A., Multivariate observations. John Wiley & Sons: 2009; Vol. 252.

61. Peterson, L. E., K-nearest neighbor. Scholarpedia 2009, 4 (2), 1883.

62. Von Luxburg, U., A tutorial on spectral clustering. Statistics and computing 2007, 17 (4), 395–416.

