## Supporting Information for "Multispectral Fingerprinting Resolves Dynamics of Nanomaterial Trafficking in Primary Endothelial Cells"

*Mitchell Gravely[1], Daniel Roxbury[1]*
[1]Department of Chemical Engineering, University of Rhode Island,
Kingston, Rhode Island, 02881, United States

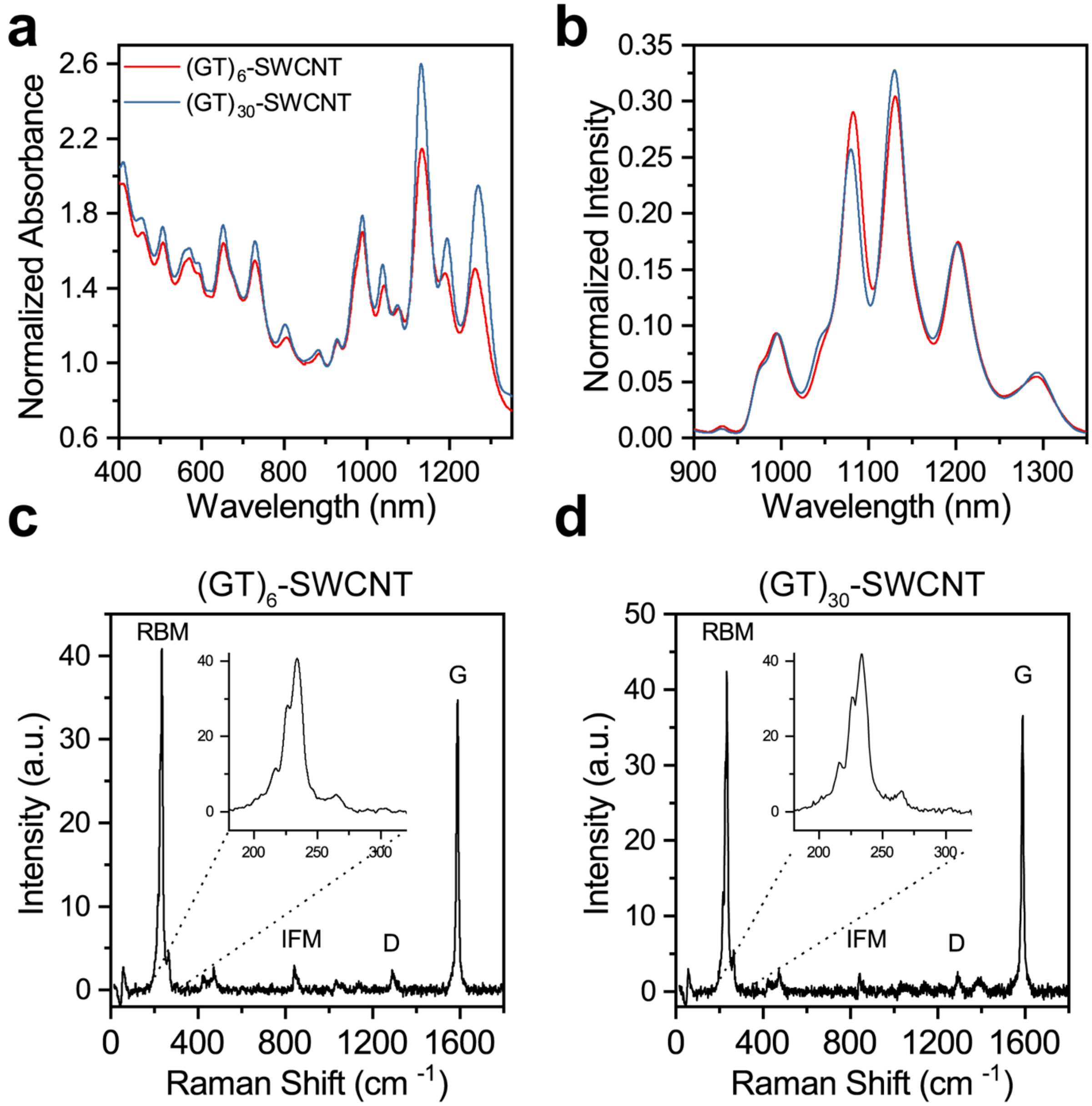


**Figure S1:** Solution-based optical characterization of DNA-SWCNTs. **(a)** Absorbance spectrum of DNA-SWCNTs in PBS, normalized to the absorbance at 910 nm. **(b)** Fluorescence spectrum of DNA-SWCNTs in cell culture media using a 730 nm excitation source, normalized to the total integrated intensity. Raman spectrum of **(c)** $(GT)_6$-SWCNTs or **(d)** $(GT)_{30}$-SWCNTs in cell culture media acquired using a 785 nm excitation source. Inset shows a close-up of the radial breathing mode (RBM) region.

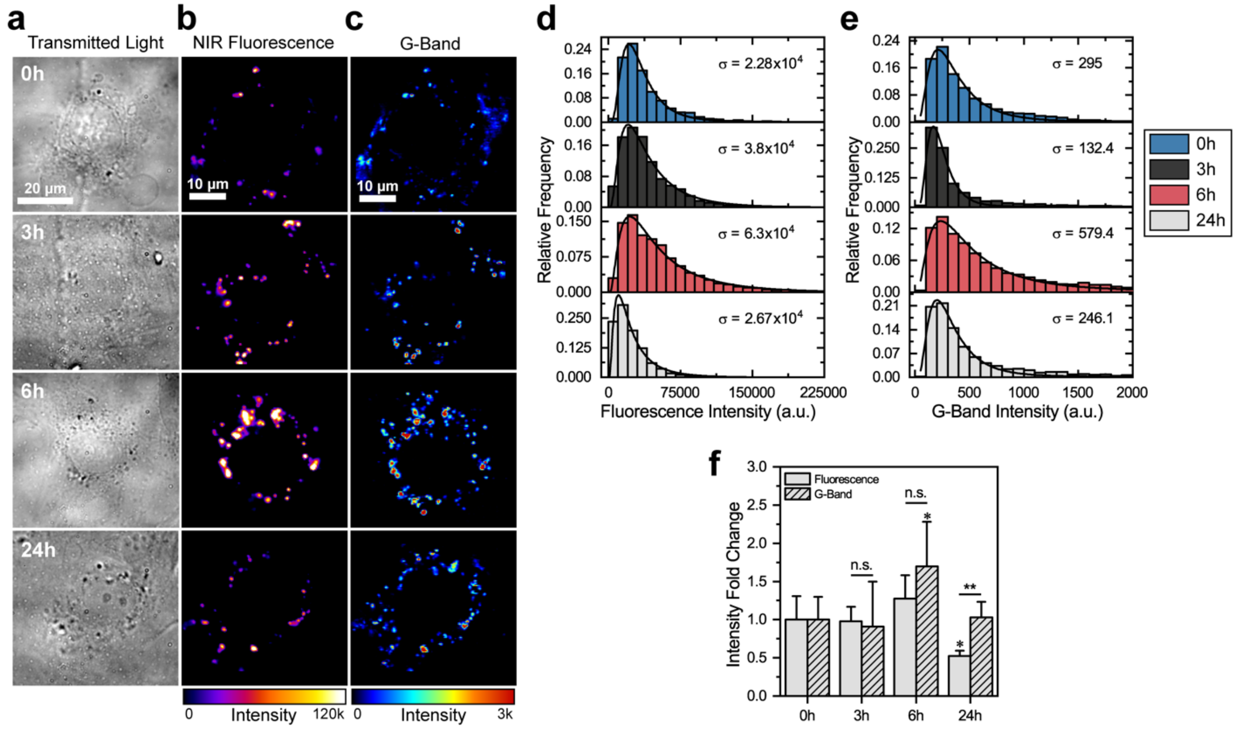


**Figure S2:** Fluorescence intensity and local concentration of DNA-SWCNTs are co-dependent within single cells. **(a)** Transmitted light, **(b)** broadband NIR fluorescence (950-1350 nm), and **(c)** G-band Raman intensity micrographs of individual cells dosed with 1 mg-L$^{-1}$ $(GT)_6$-SWCNTs for 1h and incubated in fresh media for indicated times. **(d)** Fluorescence intensity and **(e)** G-band intensity histograms of SWCNT-containing pixels from all examined cells at each time point. The distributions are fitted to log-normal curves and the widths are estimated by the log standard deviation parameter (σ). **(f)** Fold change of average fluorescence and G-band intensities with respect to 0h averages. Error bars represent mean ± s.d. with $n \geq 4$ cells per condition. Five pointed stars between columns represent significance between fluorescence and G-band intensities and six pointed stars above columns represent significance versus 0h values. (*$p < 0.05$, **$p < 0.01$, ***$p < 0.001$ according to two-tailed two-sample t-test).

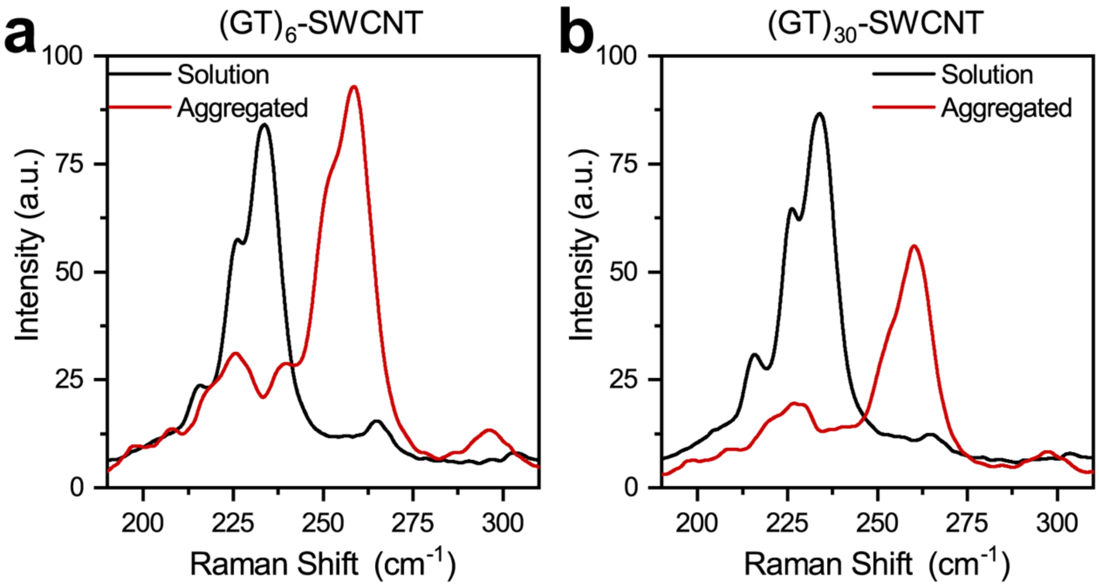


**Figure S3:** RBM of **(a)** $(GT)_6$-SWCNTs or **(b)** $(GT)_{30}$-SWCNTs in solution or intentionally aggregated and precipitated out of solution.

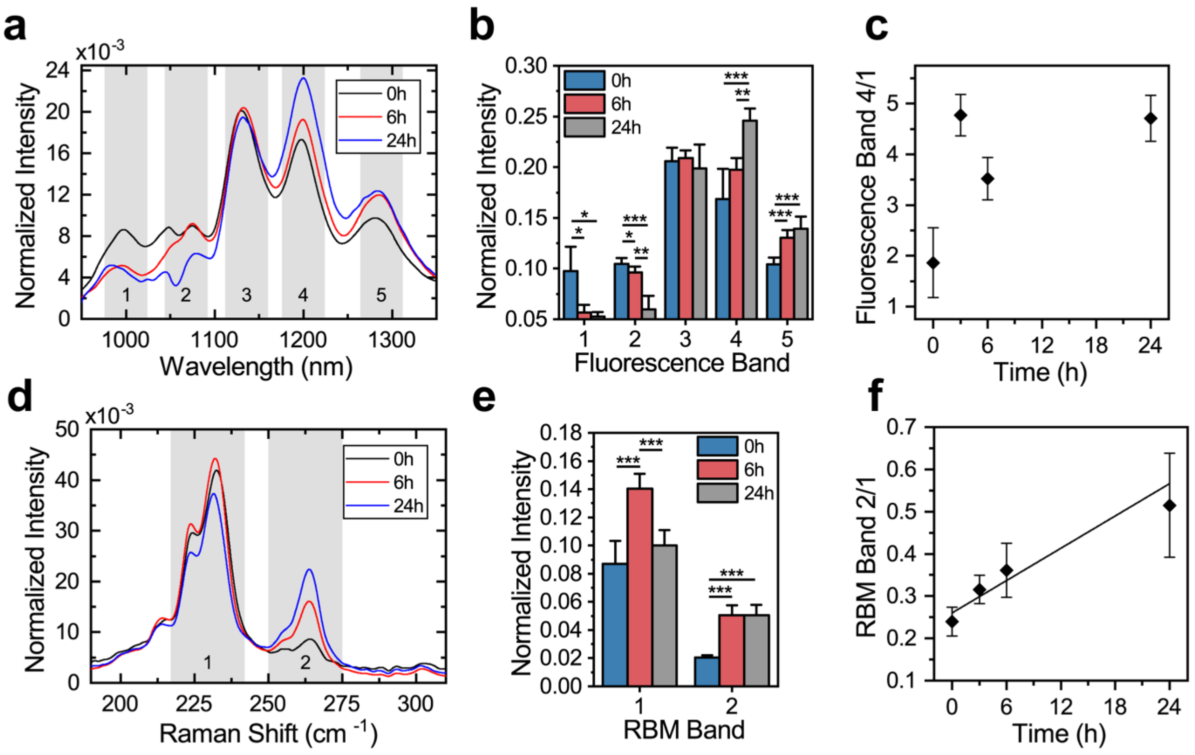


**Figure S4:** Temporal resolution of DNA-SWCNT spectral features indicates aggregation within subcellular regions. **(a)** Average fluorescence spectrum of $(GT)_6$-SWCNTs in single cells after variable lengths of intracellular processing, normalized to the total integrated intensity of each spectrum. Fluorescence bands are indicated by shaded regions. **(b)** Average normalized fluorescence band intensities from $(GT)_6$-SWCNTs in single cells after variable lengths of intracellular processing. Each spectrum was normalized by the total cell intensity, and average normalized band intensities are reported. **(c)** Ratiometric intensity of fluorescence band 4 divided by band 1 as a function of time. **(d)** RBM region of the average Raman spectrum of $(GT)_6$-SWCNTs in single cells after variable lengths of intracellular processing, normalized to the total integrated intensity of each spectrum. RBM bands are indicated by shaded regions. **(e)** Average normalized RBM band intensities from $(GT)_6$-SWCNTs in single cells after variable lengths of intracellular processing. Each spectrum was normalized by the total cell RBM intensity, and average normalized band intensities are reported. **(f)** Ratiometric intensity of RBM band 2 divided by band 1, with linear fit, as a function of time. Error bars represent mean ± s.d. for all, with $n \geq 4$ cells per condition. (*$p < 0.05$, **$p < 0.01$, ***$p < 0.001$ according to two-tailed two-sample t-test).

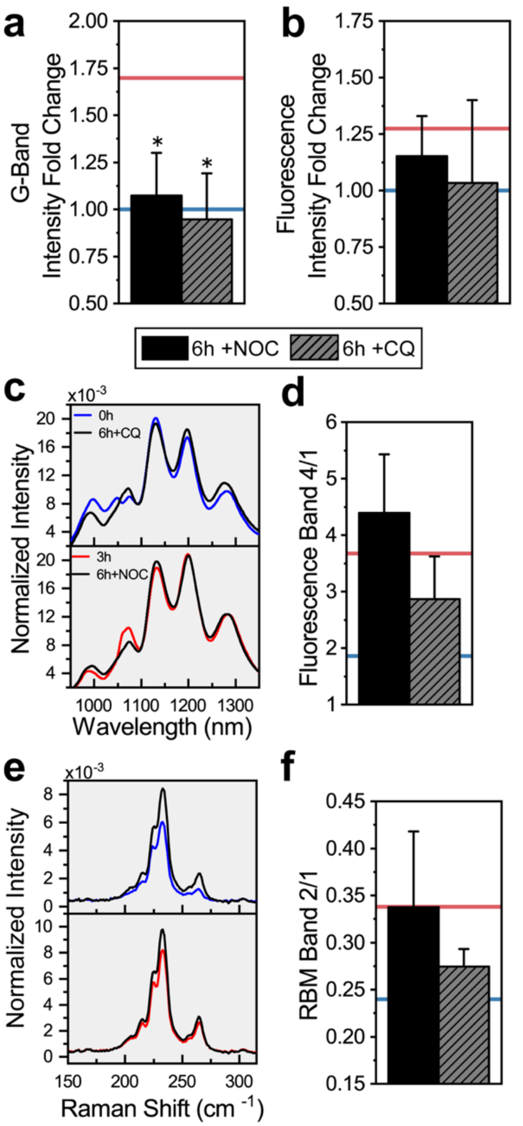


**Figure S5:** Spectral response to inhibition of endosomal progression. **(a)** Fold change of G-band and **(b)** fluorescence intensities, with respect to 0h controls, from intracellular $(GT)_6$-SWCNTs after 6h of incubation with Nocodazole (NOC, 10 µg-mL$^{-1}$) or Chloroquine (CQ, 100 µM). Averages from untreated cells at 0h or 6h are shown as blue or red lines, respectively. **(c)** Average intracellular fluorescence spectra from inhibitor-treated cells after 6h compared with fluorescence from untreated cells at indicated times, normalized to the total integrated intensity of each spectrum. **(d)** Ratiometric intensity of fluorescence band 4 divided by band 1 from inhibitor-treated cells after 6h. **(e)** Average intracellular RBM spectra from inhibitor-treated cells after 6h compared with the RBM from untreated cells at indicated times, normalized to the total integrated intensity of each spectrum. **(f)** Ratiometric intensity of RBM band 2 divided by band 1 from inhibitor-treated cells. Error bars represent mean ± s.d. for all, with $n \geq 4$ cells per condition. Stars above error bars represent significance versus 6h untreated cells. (*$p < 0.05$, **$p < 0.01$, ***$p < 0.001$ according to two-tailed two-sample t-test).

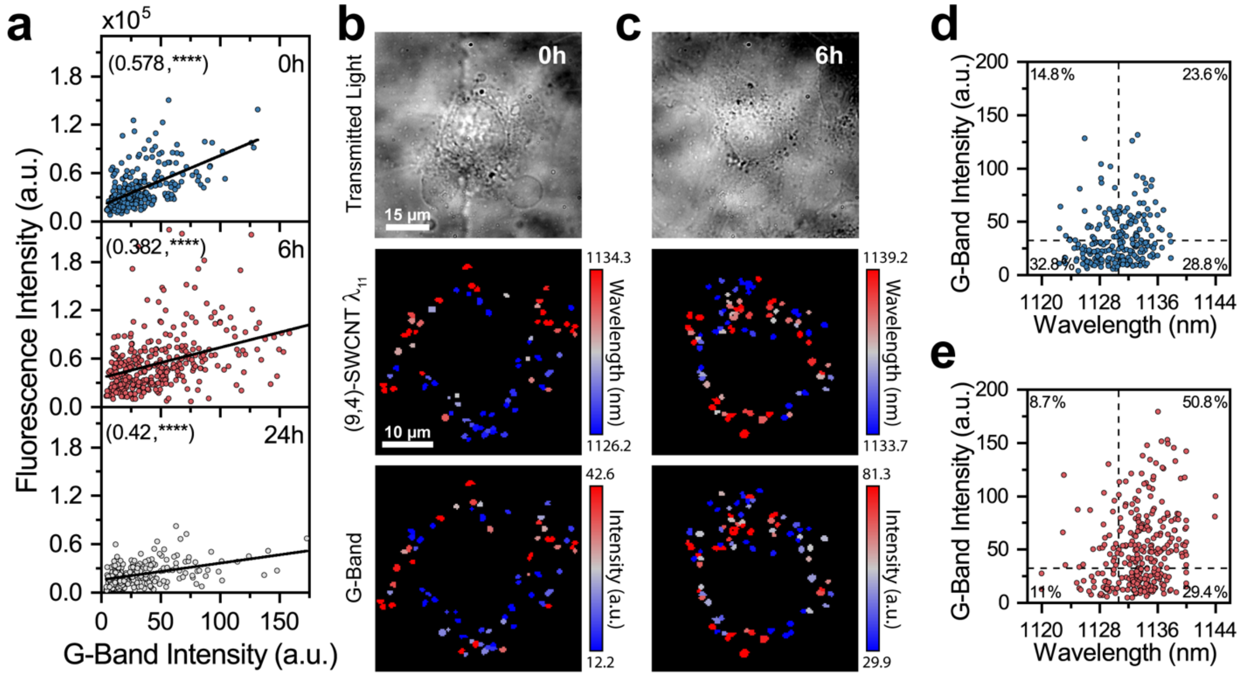


**Figure S6:** Fluorescence modulation from DNA-SWCNTs within concentrated subcellular regions. **(a)** $(GT)_6$-SWCNT fluorescence intensity as a function of G-band intensity from all intracellular ROIs, with linear fits, at indicated time points. Pearson correlation coefficients, displayed in parentheses, were calculated from scatter data at each time point. Transmitted light images, (9,4)-SWCNT emission maps, and G-band intensity maps of individual cells at **(b)** 0h or **(c)** 6h time points. Color scale range encompasses 20 – 80% of values from each ROI map. **(d)** G-band intensity as a function of (9,4)-SWCNT emission wavelength from all 0h or **(e)** 6h intracellular ROIs. Average values from 0h data, represented as dashed lines, were used to compute the percent of ROIs in each quadrant.

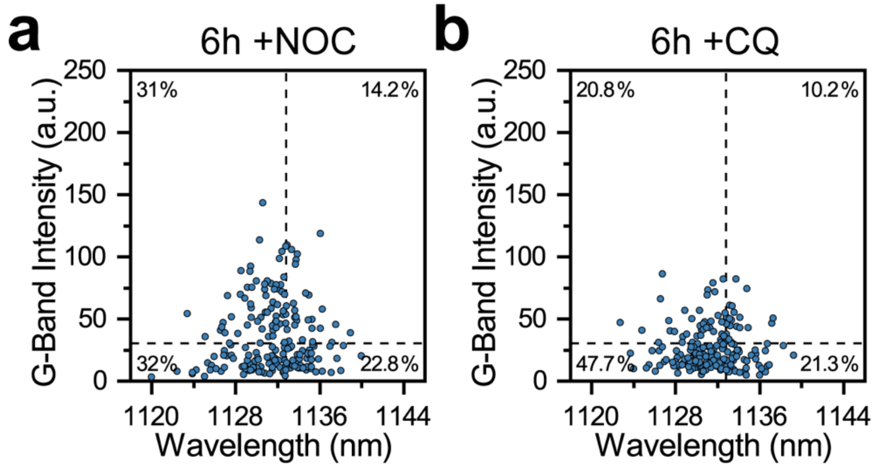


**Figure S7:** G-band intensity of $(GT)_{30}$-SWCNTs as a function of (9,4)-SWCNT emission wavelength of all ROIs from cells treated with **(a)** 10 μg-mL$^{-1}$ Nocodazole or **(b)** 100 μM Chloroquine for 6h after initial DNA-SWCNT exposure. Average values from untreated 0h cells, represented as dashed lines, were used to compute the percent of ROIs in each quadrant.

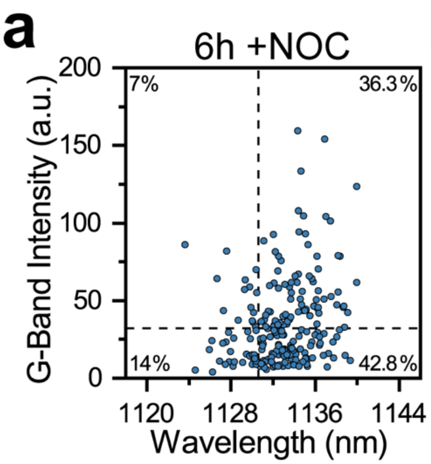


**Figure S8:** G-band intensity of $(GT)_6$-SWCNTs as a function of (9,4)-SWCNT emission wavelength of all ROIs from cells treated with **(a)** 10 μg-mL$^{-1}$ Nocodazole or **(b)** 100 μM Chloroquine for 6h after initial DNA-SWCNT exposure. Average values from untreated 0h cells, represented as dashed lines, were used to compute the percent of ROIs in each quadrant.

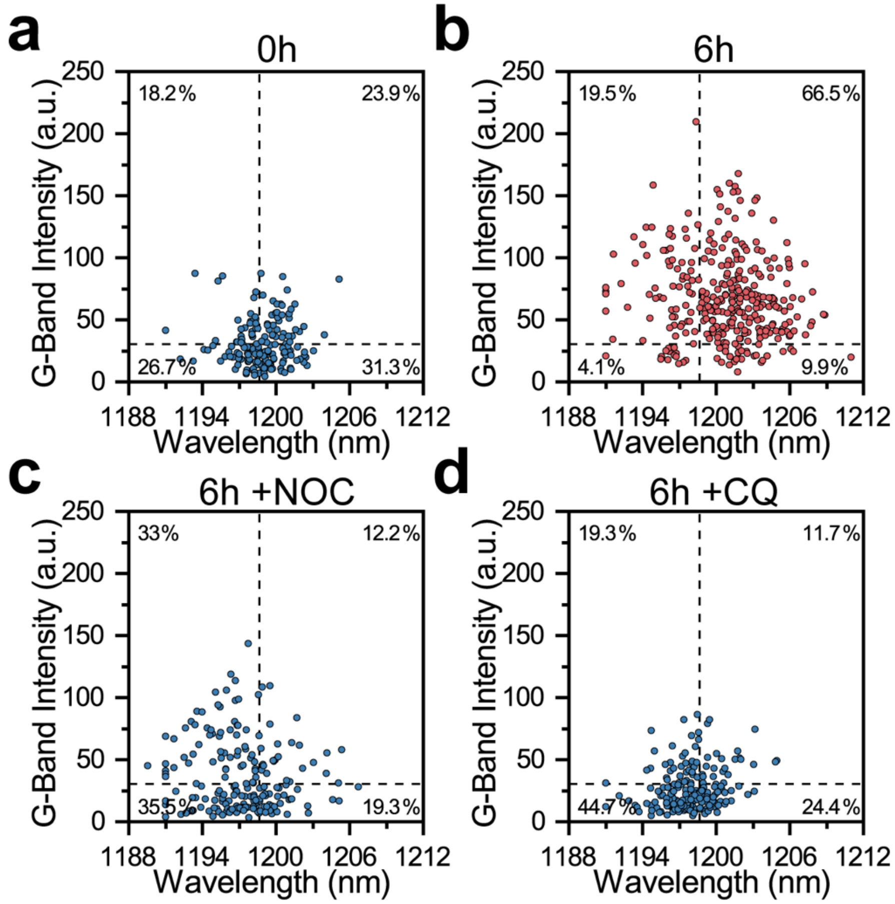


**Figure S9:** G-band intensity of $(GT)_{30}$-SWCNTs as a function of (8,6)-SWCNT emission wavelength from all **(a)** 0h or **(b)** 6h intracellular ROIs, and from cells treated with **(c)** 10 μg-mL$^{-1}$ Nocodazole or **(d)** 100 μM Chloroquine for 6h after initial DNA-SWCNT exposure. Average values from untreated 0h cells, represented as dashed lines, were used to compute the percent of ROIs in each quadrant.

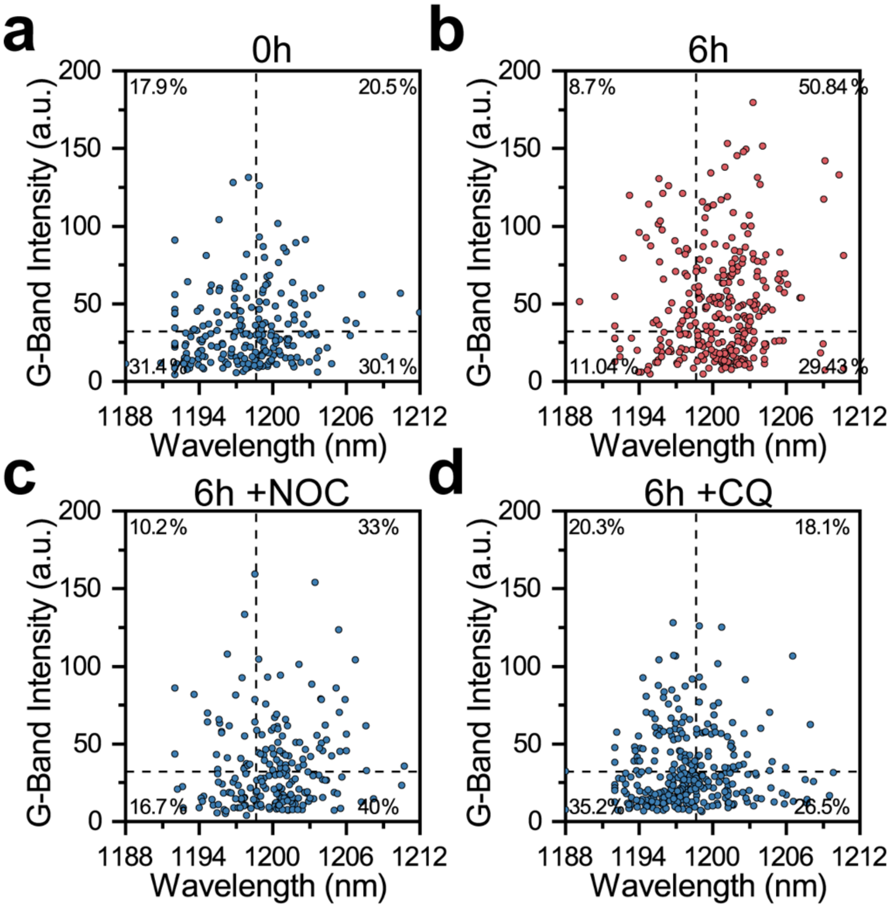


**Figure S10:** G-band intensity of $(GT)_6$-SWCNTs as a function of (8,6)-SWCNT emission wavelength from all **(a)** 0h or **(b)** 6h intracellular ROIs, and from cells treated with **(c)** 10 μg-mL$^{-1}$ Nocodazole or **(d)** 100 μM Chloroquine for 6h after initial DNA-SWCNT exposure. Average values from untreated 0h cells, represented as dashed lines, were used to compute the percent of ROIs in each quadrant.

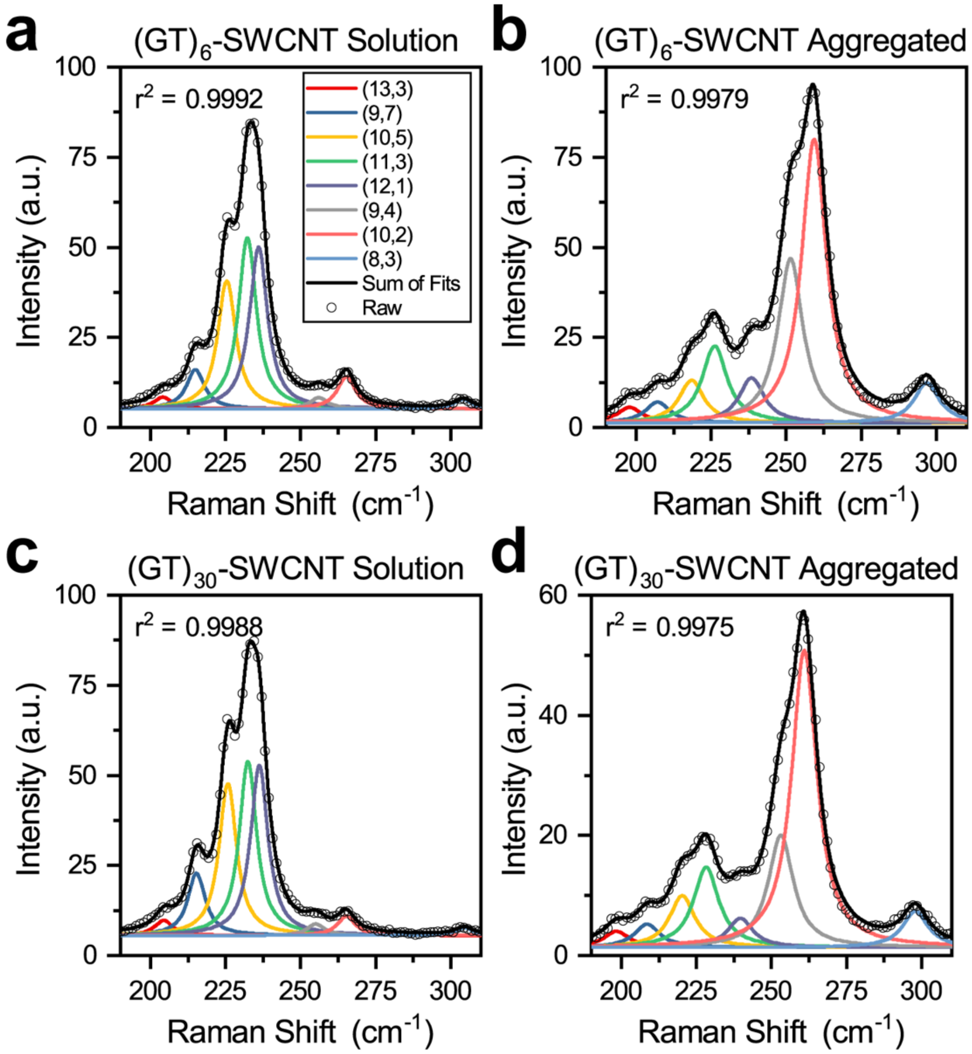


**Figure S11:** Spectral deconvolution of the RBM Raman spectrum of $(GT)_6$-SWCNTs **(a)** in solution or **(b)** aggregated and precipitated out of solution and $(GT)_{30}$-SWCNTs **(c)** in solution or **(d)** aggregated and precipitated from solution.

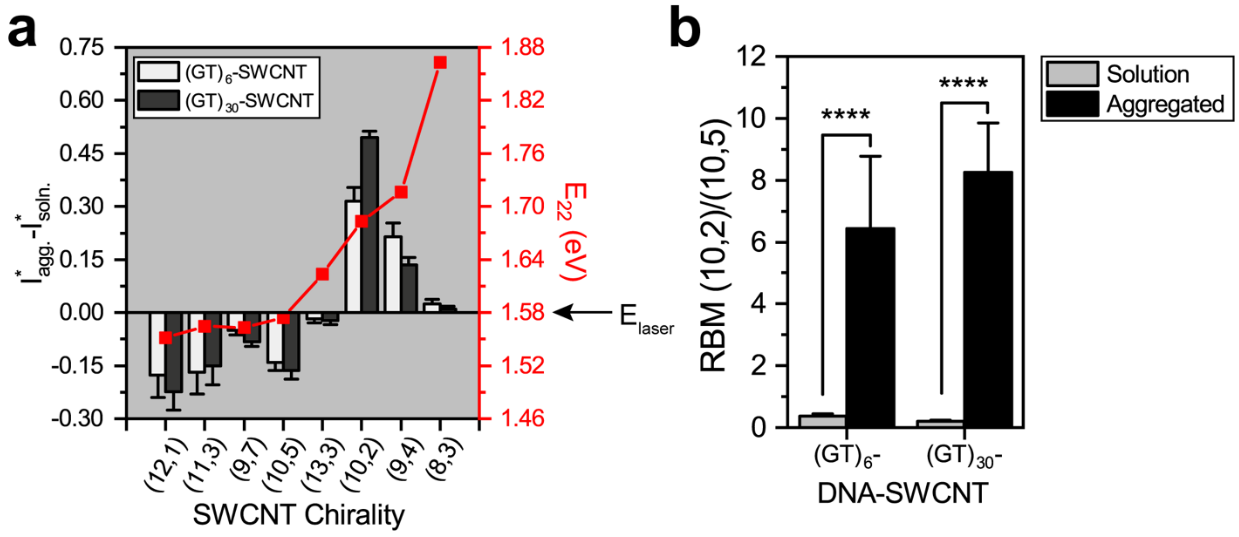


**Figure S12: (a)** Bar graph showing the relative change of RBM intensities when DNA-SWCNTs are precipitated out of solution as a function of sequence and chirality (left) with the theoretical $E_{22}$ of each chirality in surfactant dispersion overlaid (right) [1]. Each chirality intensity was normalized by the total RBM intensity from each replicate, and average intensity changes are reported. **(b)** The ratio of (10,2)/(10,5) RBM intensities from controls in solution or aggregated and precipitated out of solution. Error bars represent mean ± s.d. for all, with $n \geq 50$ spectra for each condition. (****p < 1e-4 according to two-tailed two-sample t-test).

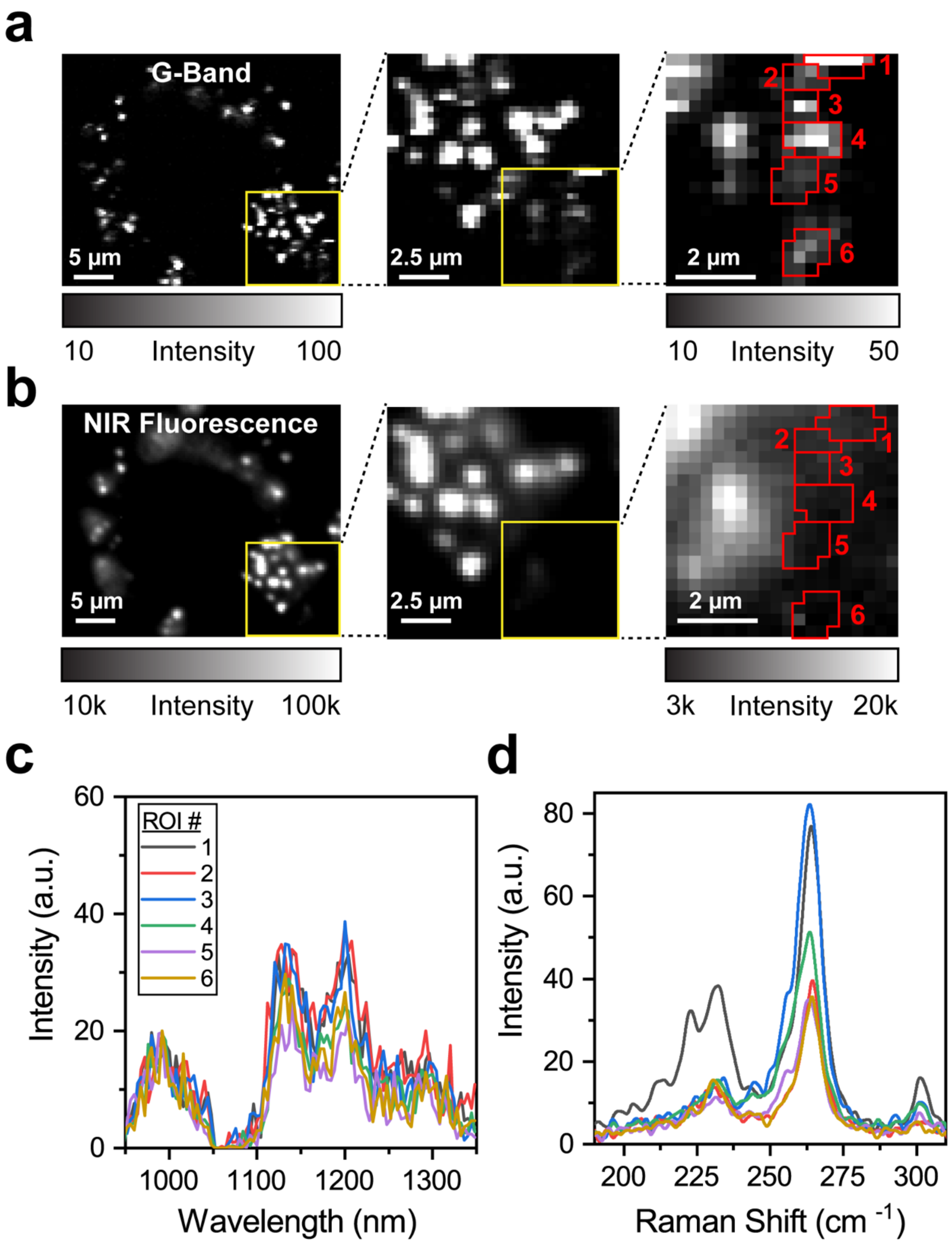


**Figure S13**: Quenched fluorescence of DNA-SWCNTs. **(a)** G-band intensity and **(b)** NIR fluorescence intensity micrographs of a single cell dosed with $(GT)_{30}$-SWCNTs and incubated for 24h. Outlined regions signify ROIs which contain DNA-SWCNTs with quenched fluorescence. **(c)** Fluorescence and **(d)** RBM spectra from the ROIs identified in (a) and (b).

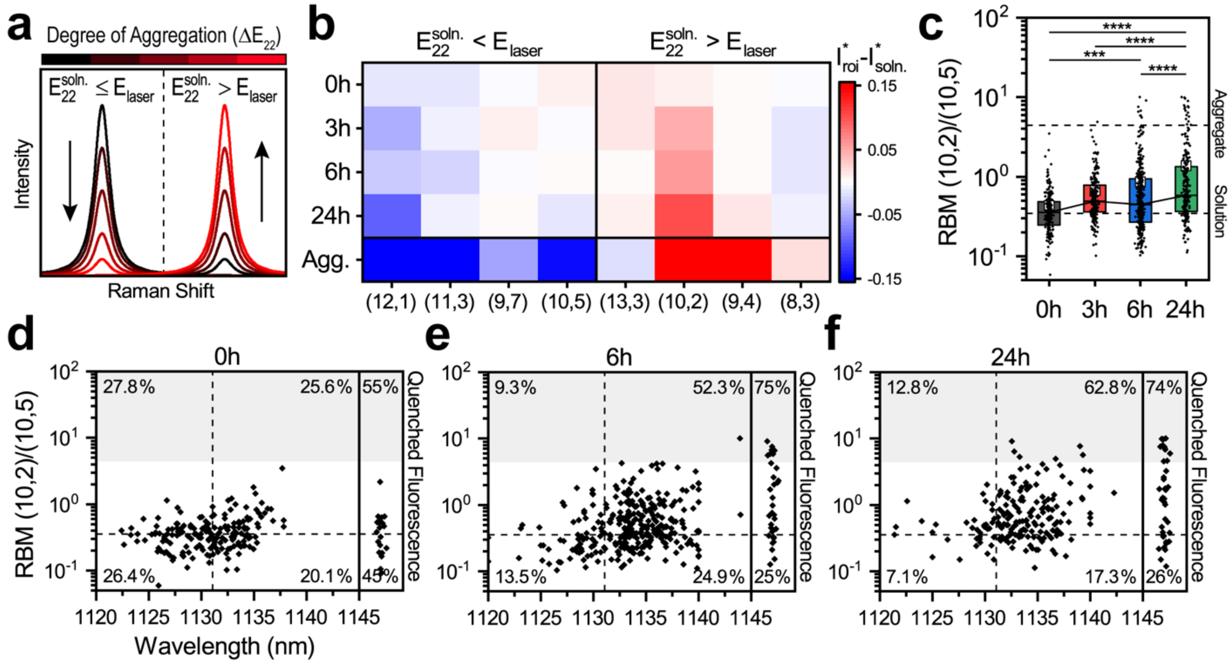


**Figure S14:** Intracellular aggregate formation is time dependent. **(a)** The RBM peak intensity of a single SWCNT depends on its transition energy ($E_{22}$) and the excitation energy ($E_{laser}$). Aggregation shifts the optical transition to lower energies ($\Delta E_{22}$), resulting in selective intensity enhancement for chiralities brought into resonance ($E_{22}^{soln} \geq E_{laser}$) and intensity reduction for chiralities brought out of resonance ($E_{22}^{soln} \leq E_{laser}$) with the excitation. **(b)** Heat map representing the change of $(GT)_6$-SWCNT RBM intensities from solution as a function of chirality and time within the cells. Control intensities of intentionally aggregated $(GT)_6$-SWCNTs are displayed as a reference. The chirality intensities from each ROI or control replicate were normalized by the total RBM intensity and average values are reported. **(c)** The ratio of (10,2)/(10,5) RBM intensities of all intracellular ROIs as a function of time. Boxes represent 25-75% of the data, small white squares represent the mean, trend lines connect medians, and dashed lines indicate values from aggregated or solution controls. One-way ANOVA with Tukey post hoc analysis was performed (*$p < 0.05$, **$p < 0.01$, ***$p < 0.001$, ****$p < 1e\text{-}4$). The ratio of (10,2)/(10,5) RBM intensities as a function of (9,4)-SWCNT emission wavelength of all **(d)** 0h, **(e)** 6h, or **(f)** 24h ROIs. Boxed column scatter plots on the right-hand side depict RBM ratio values from ROIs with poorly fitting or quenched fluorescence. Median values from 0h data, represented as dashed lines, were used to compute the percent of ROIs in each quadrant. Shaded regions indicate the (10,2)/(10,5) RBM intensity threshold identified from aggregated controls.

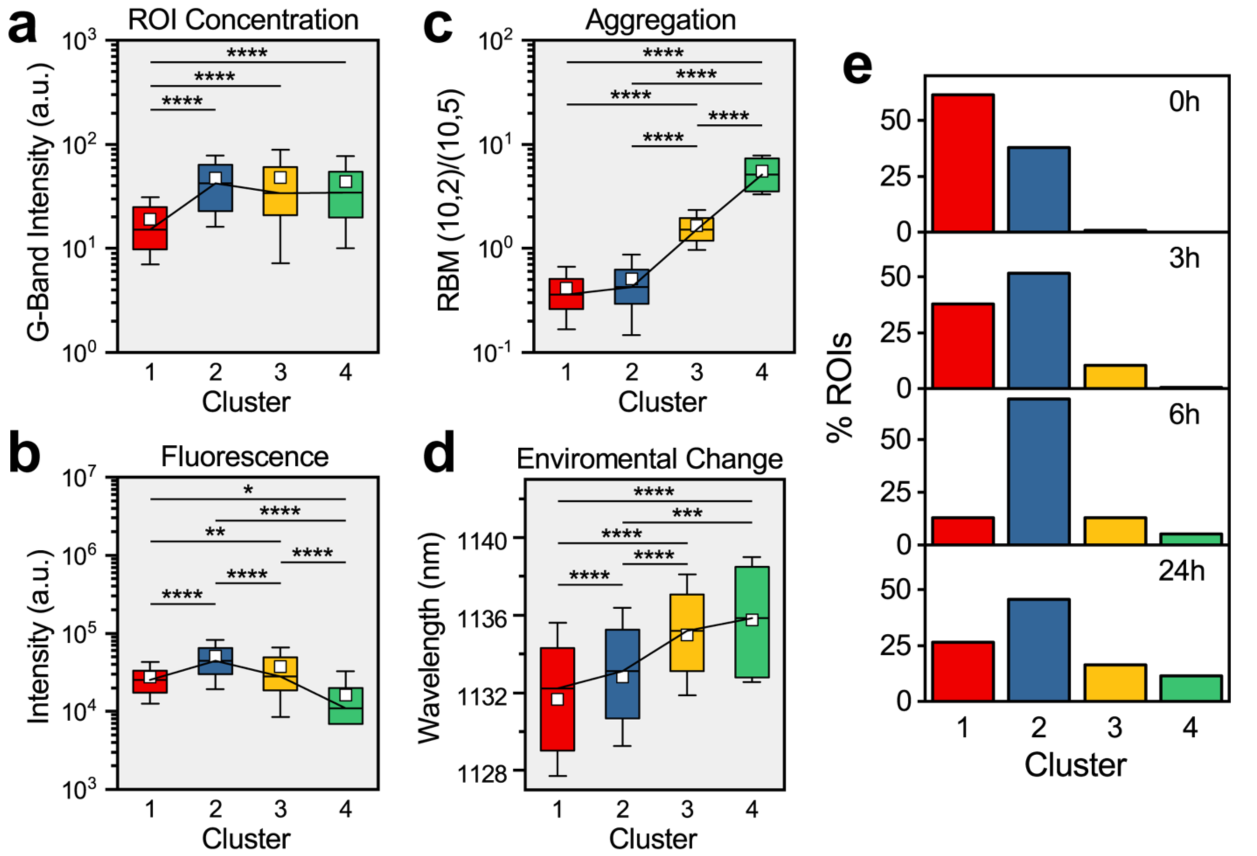


**Figure S15:** Spectral *k*-means clustering facilitates vesicle mapping *via* DNA-SWCNT spectral fingerprint. **(a)** G-band intensity, **(b)** fluorescence intensity, **(c)** RBM (10,2)/(10,5) intensity, and **(d)** (9,4)-SWCNT emission wavelength of ROIs containing $(GT)_6$-SWCNTs which were assigned to each cluster. Boxes represent 25-75% of the data, small white squares represent means, trend lines connect medians, and whiskers represent mean ± s.d. One-way ANOVA with Tukey post hoc analysis was performed for each metric. (*$p < 0.05$, **$p < 0.01$, ***$p < 0.001$, ****$p < 1e-4$). **(e)** The percent of ROIs at each time point that were classified into each cluster.

| Band | Range (nm) | SWCNT Chiralities | $\lambda_{11}$ (nm)[1] | $E_{11}$ (eV)[1] |
|---|---|---|---|---|
| 1 | 976 - 1024 | (8,3) | 968.78 | 1.280 |
| | | (6,5) | 982.36 | 1.262 |
| 2 | 1044 - 1092 | (7,5) | 1042.29 | 1.190 |
| | | (10,2) | 1074.94 | 1.154 |
| 3 | 1112 - 1160 | (8,4) | 1123.04 | 1.104 |
| | | (9,4) | 1125.14 | 1.102 |
| 4 | 1176 - 1224 | (8,6) | 1193.74 | 1.039 |
| 5 | 1264 - 1312 | (10,5) | 1275.28 | 0.972 |
| | | (8,7) | 1280.43 | 0.968 |

**Table S1:** Estimate of the optical properties of DNA-SWCNT chiralities identified in the fluorescence spectrum when excited by a 730 nm laser.

| Band | Range ($cm^{-1}$) | SWCNT Chiralities | RBM ($cm^{-1}$)[2] | $E_{22}$ (eV)[2] |
|---|---|---|---|---|
| 1 | 217 - 242 | (10,5) | 225.3 | 1.577 |
| | | (11,3) | 233 | 1.565 |
| | | (12,1) | 237.2 | 1.556 |
| 2 | 250 - 275 | (9,4) | 256.6 | 1.722 |
| | | (10,2) | 265.3 | 1.689 |

**Table S2:** Estimate of the optical properties of DNA-SWCNT chiralities identified in the RBM region of the Raman spectrum when excited by a 1.58 eV laser.

[1] Roxbury, Daniel, et al. "Cell membrane proteins modulate the carbon nanotube optical bandgap via surface charge accumulation." *ACS nano* 10.1 (2016): 499-506.
[2] Weisman, R. Bruce, and Sergei M. Bachilo. "Dependence of optical transition energies on structure for single-walled carbon nanotubes in aqueous suspension: an empirical Kataura plot." *Nano letters* 3.9 (2003): 1235-1238.
